# Ocean current models predict spatial structure for coastal biodiversity

**DOI:** 10.64898/2026.11.09.764070

**Authors:** C. Anderson Smith, Kelly N. Petersen, James M. Pringle, John P. Wares

## Abstract

Genomic diversity provides the raw material through which all evolutionary processes occur. Accurately describing how environmental features affect patterns of diversity shows promise in predicting the resilience of populations and entire communities to dynamic environments and novel stressors. Identifying the contribution of purely neutral responses to environmental effects can be difficult, especially across large spatial ranges with many covarying environmental features. Using an individual-based model of larvae carried by ocean currents, we predict patterns of neutral genomic diversity in a passively dispersed coastal species, suggesting large geographic regions of relative homogeneity with ancestry determined by the nearest “upstream” population with limited larval input. The genomic identity that dominates each region sharply transitions at the next such interruption, generating hierarchical structure and violating assumptions of isolation-by-distance. We call these upstream populations “lineage source regions” (LSRs), and we find strong concordance between LSR model predictions and an empirical population genomic survey of a keystone salt marsh mussel (*Geukensia demissa*). We quantitatively validate model performance and goodness-of-fit, updating baseline predictions of genomic diversity patterns from the classic isolation-by-distance model to this LSR framework. This spatially explicit coastal ancestry model has potential for generalization across passive dispersing species.

**Significance Statement:** We construct an individual-based ocean current model to simulate coastal population genetic structure of a passively-dispersed organism, and collect empirical genomic data from a wide-spread coastal mussel for comparison. We quantitatively demonstrate the superiority of this approach as compared to the typical assumption that isolation-by-distance generates and structures neutral genetic diversity in the ocean. This paper proposes an explicit environmental mechanism behind neutral population genomic structure across spatial samples, allowing improved accuracy in characterization of neutral patterns of diversity before attempting to pinpoint adaptive signals.

## Introduction

Genomic diversity not only serves as a foundation for population persistence (1), but patterns of genomic diversity and divergence have the potential to influence entire communities, alter the function of ecosystems, and predict local resilience against stressors including environmental change (2–6). Anticipating such responses requires accurate characterization of the specific evolutionary mechanisms that determine how patterns of genomic diversity emerge, change, and persist (7, 8).

Population genomic approaches to characterize how genomic diversity is moved and maintained in natural systems vary widely, with some screening for strong adaptation as an *a priori* mechanism and others focusing on neutral demographic fluctuations across time and space (9–12). Natural selection exerts strong, environmentally mediated effects both on particular genomic regions and quantitative traits (13). However, when population genomic surveys focus first on adaptive explanations for observed patterns of diversity, they often generate a compelling yet incomplete description of the focal organism’s interaction with its environment (14–16). In spatial or temporal samples of genomic diversity, patterns of adaptation can only be fully characterized after describing how organisms and populations respond to neutral mechanisms such as movement or dispersal because neutral genetic structure itself skews tests of adaptation (14, 17–19).

Often, the null hypothesis for neutral divergence is described by movement models developed decades ago; we assume isolation-by distance (IBD) as a basic but informative mechanism driving population genetic structure across spatial samples (20). Yet more accurate neutral null models can be parameterized to account for environmental features, demography, or evolutionary history (21–23). Ocean currents are one such environmental process hypothesized to structure genomic diversity in a wide variety of taxa by influencing how offspring move between patches of suitable habitat (24–27). Many marine species’ distributions of genomic diversity are likely structured by isolation by resistance (IBR), where the dispersal probability given known current patterns may better predict genetic relatedness than spatial distance alone (21, 24, 25, 27, 28). However, this relationship may be more complicated than is currently accepted. Across a species’ distribution, diverse coastal features – spatial variation in upwelling, winds, and buoyancy forcing, along with changes in shelf depth, can create discontinuities in flows of larvae which serve as discontinuities in gene flow (29).

The protected areas that form immediately downstream from these interruptions in larval delivery (30–33) may have strong impacts on population structure by forming and maintaining sources of genotypes at the sites of interruptions that themselves seed larger downstream areas of relative genetic homogeneity (30, 34, 35). We refer to these areas as lineage source regions (LSRs), and we predict that they will have higher persistence of genotypes that will subsequently swamp downstream sites, homogenizing the genetic diversity of a broader region (31).

We hypothesize that variation in the prevalence of genotypes at a location is the result of metapopulation dynamics among genomic lineages, where the migration rate of genotypes to a location shapes genomic composition across space without requiring differential fitness between genotypes (34, 36, 37). The numerical advantage possessed by propagules from LSRs renders downstream regions “sinks” of genomic diversity (31, 34). This would result in hierarchical structure as well as a homogenization of genomic diversity between each interruption in larval delivery (36, 37). Because ocean currents function as larval transporters across space, the LSR model can be generalized to a variety of passively dispersed organisms, such as grain pests dispersing along human shipping routes (38), wind-dispersed fungal spores (39), and riverine species (40).

The ribbed mussel *Geukensia demissa* is an ideal organism to test these predictions of the LSR framework due to its passively dispersing larvae and well-characterized life history (41–43). It is also ecologically important as its interactions with marsh cordgrass aid in salt marsh ecosystem resilience, and *G. demissa* is distributed across several major ecoregions (44, 45), across which genomic and inter-species range boundaries have been documented in other marine taxa (32, 45–47). As such, genomic diversity within species with drifting larvae are likely to be strongly affected by ocean processes and may impact the structure and function of coastal communities, providing motivation for applying the LSR model.

We use high-resolution physical ocean models for identifying the likely recent ancestry of organisms with these larval dispersal traits, thus generating expectations of neutral genetic structure. We test the LSR framework as a predictive model for passively-dispersed coastal diversity using empirical data from *G. demissa* across its range at the highest spatial and genomic resolution to date. We expect that accounting for ocean current processes as the mechanism that spatially structures genomic diversity will improve the explanatory power of neutral models over simpler models that assume isolation-by-distance, which may permit more precise characterization of genomic diversity that responds via adaptation to environmental gradients.

## Results

### Population genomics

We sampled *Geukensia demissa* from 23 coastal locations between Wakulla Beach, Florida and Antigonish, Nova Scotia (Supplementary Table 1) and prepared libraries using the 3RAD protocol illustrated in (48). Sequence data for 224 individuals, assembled *de novo* in STACKS, contained an average of 7.16 million reads per individual; all data are at NCBI BioProject PRJNA977713. These data include individuals representing the congeneric *G. granosissima* from the Gulf of Mexico. Following (49), our sample at Lake Worth, Florida is excluded from analysis here because of historic introgression patterns between the two species. We also excluded 26 individuals due to high values of missing data or evidence of introgression with the Gulf Ribbed mussel (details in Supplemental Information Part B). The individuals removed due to evidence of introgression originated from Roanoke Sound, North Carolina Gloucester Point, Virginia. Filtering and outlier screening of the final sample set yielded 9249 loci.

To determine the genomic structure across the range, we utilized two *k-*means clustering algorithms, sparse non-negative matrix factorization (sNMF) and discriminant analysis of principal components (DAPC). Results from DAPC delineated 5 clusters (See Supplementary Figure 14). Samples from the Canadian Maritimes differentiate from Gulf of Maine samples along PC2, and both groups differentiate from the rest of the sampling distribution along PC1. The lowest cross-entropy values following sparse non-negative matrix factorization (sNMF) suggest values of *k* ancestral populations between 2 and 4 are most appropriate. We report all three clustering distributions in Supplementary Figure 16 and compare LSR predictions to *k = 4.* At *k=2,* sNMF delineates a southern and northern genotype class that transitions rapidly at Cape Hatteras, followed by a genotype class corresponding to samples from the Canadian Maritimes at *k = 3* and separating the Gulf of Maine at *k = 4* (Figure 1). The same spatial structure is present with or without exclusion of outlier loci.

**Figure 1.**
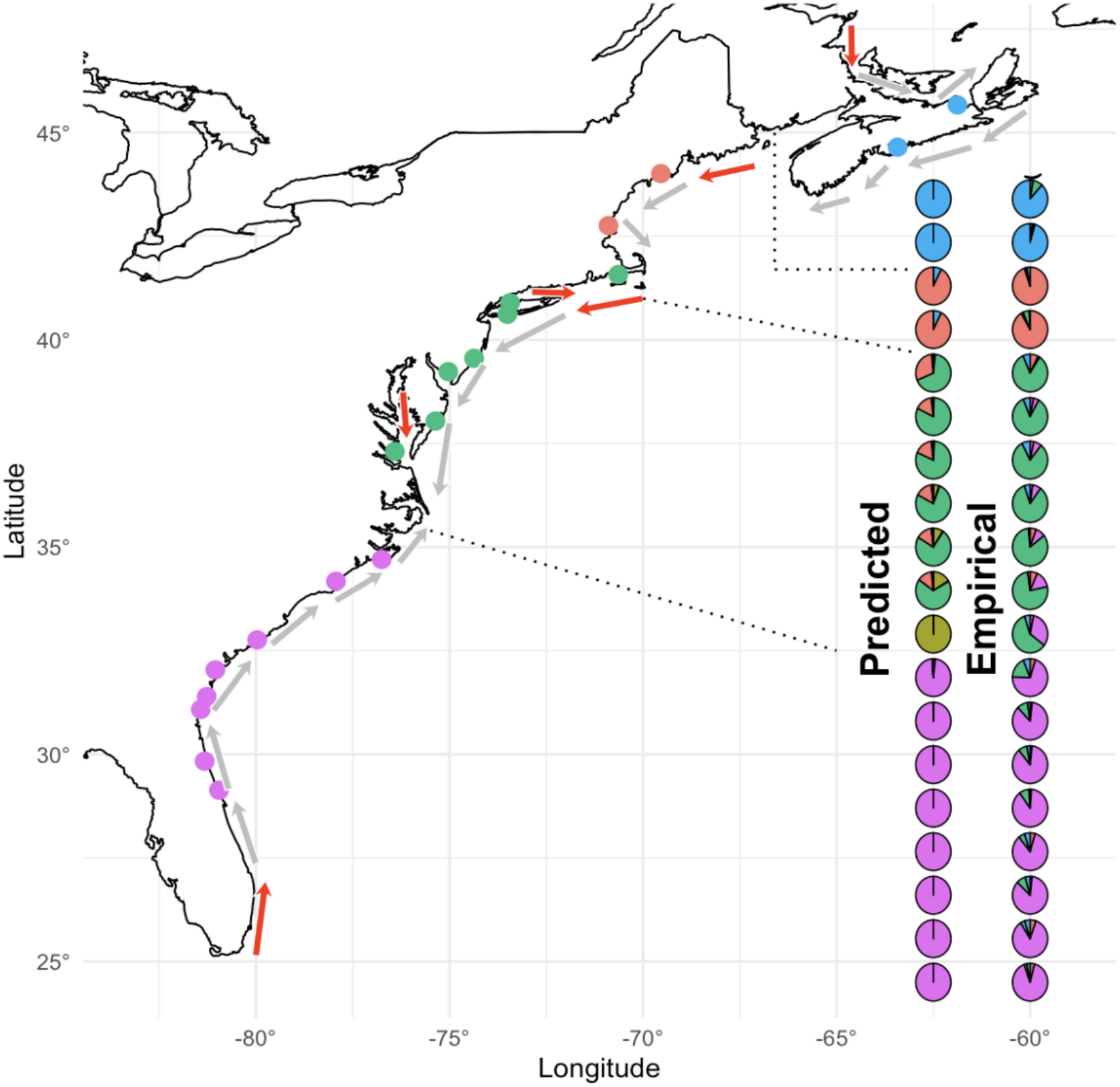
Map of sampling locations and ocean current patterns, where point colors represent dominant snmf-assigned ancestry group and arrows represent ocean current flows, with red arrows representing flows that generate LSRs. Pie chart colors represent predicted genomic ancestry via the LSR model (“Predicted”) and mean snmf-assigned ancestry group for G. demissa (“Empirical”). Dotted lines represent breaks in dominant lineages predicted by LSR framework as well as boundaries between major ecoregions (in order from south to north: pink points represent the South Atlantic Bight, green points represent the Mid-Atlantic Bight, red points represent the Gulf of Maine, and blue points represent the Canadian Maritimes).

To screen for isolation-by-distance across the range, we applied a variant of matrix correlation test known as the Mantel test to pairwise genomic and geographic distance matrices of samples from each location.The Mantel test on pairwise linearized F_ST_ and geographic distance for the entire range of the *G. demissa-*only dataset suggests isolation by distance (r = 0.6827, p = 0.001), however this pattern is not replicated within regions delimited by the above clustering analyses, indicating high relative homogeneity within these regions. Such tests were not possible within the Gulf of Maine or Canadian Maritimes due to low sampling density, but neither the Mid-Atlantic Bight (r = 0.3058, p = 0.154) nor the South Atlantic Bight ( r = 0.1367, p = 0.278) regions displayed any signal of isolation-by-distance despite a geographic extent of over 1000 kilometers. Pairwise F_ST_ values are elevated between LSR-driven areas and depressed within them, with the Canadian Maritimes (Antigonish and Halifax, NS) and the Gulf of Maine (Damariscotta, ME and Plum Island Ecosystem, MA) displaying the strongest signal of fixation from the rest of the range (Figure 2).

**Figure 2.**
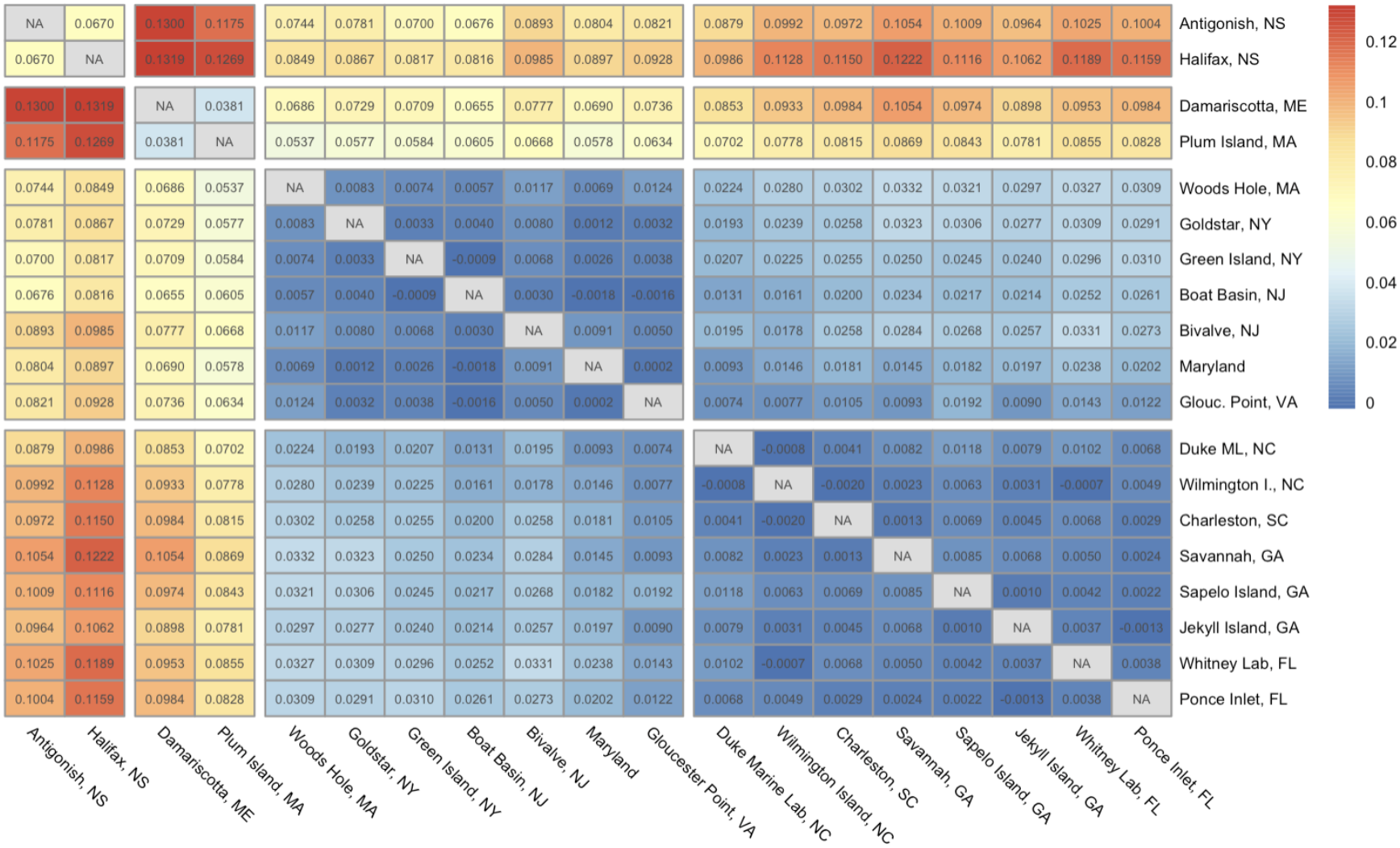
Heatmap displaying pairwise Weir-Cockham F_ST_ between sampling sites, arranged from north-south left to right and top to bottom. Gaps between groups of cells indicate a shared predicted LSR between them; groups within these gaps correspond to marine ecoregions.

### Current modeling

We generated an individual-based model to predict larval dispersal and thus genomic identity based on the flow of ocean currents. A neutral prediction of the spatial genetic diversity resulting from ocean current-driven dispersal followed a numerical model parameterized by larval life history of *G. demissa*. When targeting an 18-day larval duration to match experimental assessment of pelagic duration (50) using global physical ocean data from 2007 to 2024, modeled ancestry proportions indicate individuals living along large portions of coastline descended from ancestors concentrated in a few very localized regions (LSRs denoted with black arrows in Figure 1). Modeled individuals throughout the range of *G. demissa* can be characterized by the relative proportion of the individual’s lineage that descend from each distinct LSR, and there are sharp geographical breaks in these proportions. The model leads to consistent predictive outcomes if the larval duration is decreased or increased by 8 days (Supplemental Information Part H, Supplementary Table 10).

### Comparative oceanographic-genomic analysis

To model the relationship between empirical genomic identity and the predictions of the LSR model, we utilized redundancy analysis (RDA), a method of genotype-environment association (GEA) generally preferred for its low false positive rate (51). The RDA models of sampled genotypes relative to predicted LSR ancestry, temperature, and latitude were all highly significant relative to null variables (p < 0.001). Of these results, the LSR-ancestry model explained the most variation (r-squared = 0.56, adjusted r-squared = 0.28), followed by temperature (adjusted r-squared = .11 and latitude (adjusted r-squared = 0.11). In the LSR model, RDA1 captured 16.94 percent of all variation, and RDA2 captured 11 percent. A semivariogram of the LSR model residuals from RDA1 indicated no spatial structure among residuals (Supplementary Figure 18).

Goodness of fit to our empirical genomic dataset, assessed by calculating Hellinger distances across all sites, was higher for the LSR model than for an IBD simulation generated using SLiM (Figure 3). The outcomes for modeled PLD of 10 or 18 days have a nearly equivalent fit to the genomic data, and are in the 94th percentile relative to the low-migration simulations of isolation by distance, 89th percentile relative to the middle migration value, and 91st percentile relative to the high-migration IBD simulations (Figure 3). The LSR model with PLD of 26 days is still in the upper quartile relative to all IBD simulations (Figure 3). The greatest potential overlap among LSR-driven regions is between the two predicted areas within the Mid-Atlantic Bight (MAB; between Cape Hatteras and Cape Cod) and between the northern MAB predicted region and that of the Gulf of Maine. We present the results combining the two MAB predicted regions.

**Figure 3.**
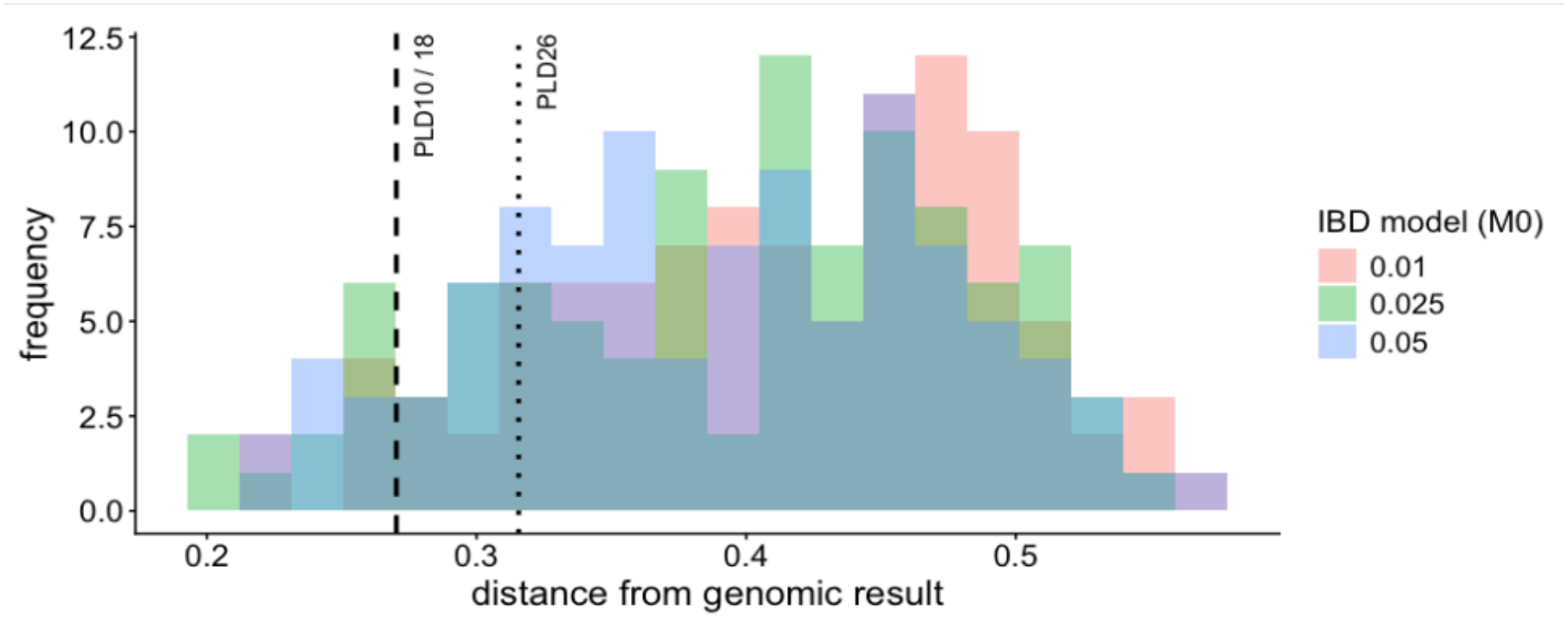
Histogram of Hellinger distances from observed sNMF-estimated ancestry for K=4 relative to simulated isolation by distance data for the same spatial samples. Hellinger’s distances to the best-fit summaries of the “ocean” model indicated with a single line for pelagic larval duration of 10 and 18 days, as they overlap.

## Discussion

Population genomic data from *G. demissa* indicate the presence of strong hierarchical structure, with cluster boundaries corresponding to interruptions in the continuity of larval delivery as predicted from neutral ocean dispersal patterns. Though there is also considerable environmental variation along this coast, adaptation does not need to be invoked in studying the maintenance of this pattern. The concordance between model and data is supported by high goodness of fit relative to simulated isolation-by-distance. A similar pattern of hierarchical genomic structure exists in a diverse array of taxa on the Atlantic Coast of the United States and Canada, indicating that the predictions of the LSR model can serve as an updated neutral null hypothesis for the mechanism structuring genomic diversity along coastlines, with broad generalizability to other passively dispersed organisms.

### Population genomics

Genetic identity of *G. demissa* individuals across the range is strongly predicted by the identity of the closest upstream LSR identified by our model. This finding validates the explicit role of ocean currents and the role of neutral mechanisms in the creation of hierarchical population structure in marine species with drifting larvae. Empirical clustering of ribbed mussel genotypes is more consistent with predictions from ocean current modeling than with a lack of structure, structure driven by simple isolation by distance, or a relationship with local sea surface temperature (itself a function of both latitude and ocean currents). Clustering analyses, pairwise F_ST_ values, and oceanographic modeling all pinpoint the boundaries of LSR-driven regional homogeneity in the same places: the Bay of Fundy, Cape Cod, and Cape Hatteras. Yet these canonical ecoregion boundaries are not themselves mechanisms (44, 45); our ocean current model provides that mechanism in a manner robust to a wide range of simulated larval dispersal durations.

Even though traditional approaches fail to reject isolation by distance across the range, divergence of samples within LSR-driven regions are universally low despite distance (52), and the asymmetrical nature of current-mediated transport violates the assumptions of statistical tests for IBD (22, 53). Instead, we see evidence of spatial homogeneity within LSR-driven regions and hierarchical divergence between them. Current background expectations of diversity patterns in passive dispersers predict lower clustering and a much more gradual transition of genomic identity across space in a neutral SNP dataset (54, 55). Thus, the large swaths of relative homogeneity punctuated by swift transitions to different genomic lineages at each successive interruption in larval flow are discordant with the typical null model of IBD.

Genotype frequencies vary most sharply when comparing once-glaciated regions with those to the south. Clustering analyses and comparisons of pairwise F_ST_ values show strong differentiation between the Canadian Maritimes, the Gulf of Maine, and the rest of the range (Fig. 2). Mitochondrial sequence diversity is indicative of founder events as well (56). The genotypes sampled from 19 locations along the Atlantic coast cluster cleanly in regions along the coast that the LSR model predicted would be dominated by ancestry from source regions. This pattern is driven purely by ocean physics; neither strict isolation nor adaptation are necessary for the observed spatial structure. In fact, there was no visible geographic pattern observed in potentially adaptive outlier loci identified (and removed) from our analyses, supporting the hypothesis that neutral mechanisms drive patterns of divergence described here.

Along with population structure evident along the Atlantic coast, our data captured a surprising instance of introduced diversity from the Gulf of Mexico. Introgressed mussels far from the native range of the Gulf ribbed mussel *G. granosissima* are contained in the Chesapeake Bay and Roanoke Sound samples; this congener may have been introduced accidentally during restoration of the Chesapeake Bay oyster fishery with oysters from the Gulf coast in the mid-twentieth century (57). Limitation of *G. granosissima* introgression to this region is unexpected given both species’ high dispersal potential, but remarkably, the samples are placed within yet another region predicted to be comprised of genotypes from LSRs (Figure 1, “Predicted”). Though their nuclear genome is dominated by Gulf diversity, these individuals carry mitochondrial haplotypes belonging to the Atlantic ribbed mussel ((56); see Supplemental Information Part B, Supplementary Figures 10-11).

### Current modeling

Ocean current modeling suggests that the mechanism generating and maintaining population genetic structure in this system is spatially-dependent variance in larval delivery, in which neutral allelic diversity that arises has a varying potential for persistence depending on where in a domain it arises (31, 36). Domains with interruptions in current-mediated larval input immediately preceding them will contain genotypes that persist, whereas genotypes from downstream domains will not. This punctuated asymmetric gene flow on ocean currents has the potential to dramatically lower the effective population size, compressing the coalescent time scale within these regions relative to the time scale across all sample sites (58).

The results of the physical modeling are not greatly sensitive to moderate changes in the planktonic larval duration, with similar predicted results for 10 and 26 day larval durations. The results of the model are robust to the planktonic duration because the LSRs are formed in regions where the alongshore transport, or interruption in alongshore habitat – as in the Bay of Fundy – shelters the region from migrants from elsewhere (30, 33, 35) no matter their dispersal potential. In general, a change in pelagic larval duration (PLD) will alter the magnitude of larval dispersal, but not the direction or spatial pattern (59), and thus a change in PLD will not disrupt the spatial patterns of dispersal which lead to LSRs. Unlike related approaches (27, 60) that focus on how connected pairs of samples are by ocean currents, our approach focuses on the maintenance of distinct geographical origins of the genetic lineages found in different locations.

The recent improvements in available physical ocean models have also been used to test the relationship between ocean current patterns and the establishment of introduced species (29, 33, 61). These same models can predict patterns of genomic homogeneity that stabilize across hundreds of generations or more at a more granular spatial resolution than other, vector-based frameworks or parentage models that only refer to the most recent generational structure (24, 62). Partnered with contemporary genomic approaches that generate thousands of variable loci, we can identify important spatial structure despite very small and limited range of mean pairwise fixation metrics (i.e. F_ST_; (27, 63)). Thus, the broader model-focused spatial clustering of ancestral lineages across a distribution may be a more powerful way to show concordance between model predictions and the demographic history of a coastal species.

### Comparative oceanographic-genomic analysis

Our results verify a strong explanatory relationship between oceanographic modeling and the genomic patterns described in *G. demissa* that surpasses isolation-by-distance and does not require adaptive causes. Further, we identify a distance-based approach to explicitly compare the goodness of fit of empirical population genomic data with simulated distributions under isolation-by-distance and the LSR model.

The LSR model makes predictions at a higher spatial resolution than we sampled, predicting more LSRs contributing to the ‘seascape’ of diversity on this coast than are apparent in the genotypic data; this disparity in the predicted number of LSRs relative to the number of empirical clusters penalizes the LSR model in comparison with IBD. Its strong performance despite this penalty indicates that the LSR model may be more useful in predicting patterns of coastal diversity as a neutral null model than IBD, which assumes symmetric stepping-stone dispersal.

Additionally, discrepancies in the goodness-of-fit comparison clarify where the LSR model might decouple from biological observations as well as how null IBD models often generated predictions far removed from empirical data. Model goodness-of-fit consistently deteriorates near large estuaries and sounds - with sharper boundaries in the empirical data than the model predicts - and so mechanistic explanations for this deviation will be the focus for future work.

Previous descriptions of population connectivity in other species across the Atlantic coast of North America mirror the strong hierarchical structure demonstrated in this study, where relative homogeneity exists within large regions with stark differentiation between them. At Cape Hatteras – the most prominent genomic boundary captured in *G. demissa –* species such as silverside (*Menidia menidia)*, shovel bugs (*Haustorius canadensis)*, and marsh cordgrass (*Spartina alterniflora)* experience rapid, pronounced transitions in genomic diversity (28, 64, 65). The Gulf of Maine functions as a site of genomic transition in a wide range of taxa as well, one that encompasses green crabs (66), oyster drills (67), polychaetes (68, 69), sea stars (70), cordgrass (71), and - a most direct comparison - oysters (57). At more northern latitudes, the Scotian Shelf also serves as a site of transition in organisms such as sand lances (72) and sea scallops (73).

These large stretches of homogeneity between transition points have been detected with respect to species range distributions and described as “ecoregions” in previous work (44), and their origins remain the subject of some debate (45, 74). However, the current-based LSR model presented here predicts spatial patterns of empirical genomic data, indicating that the highly consistent genomic and species transitions described by ecoregions and comparable genomic studies (47) reflects the presence of LSRs and the regional homogeneity they generate. Global applications of LSR simulation provide evidence for this relationship across taxa; the LSR model accurately predicts species ranges in concordance with empirical marine ecosystem boundaries (75, 29).

### Implications

The physical model described here – using high-resolution ocean current data and parameters based on known life history – predicts genomic hierarchical spatial structure in *G. demissa.* We expect other organisms on the Atlantic Coast of North America, especially those with planktonic dispersal, to follow this pattern. Our methods represent a novel quantitative approach with which to explain the origin and maintenance of genetic structure along coastlines, and our results indicate that ocean currents can structure species with passively-dispersing larvae through wholly neutral dynamics (29, 30, 41). The LSR framework provides an updated neutral null expectation in passively dispersed species to more accurately reflect mechanistic environmental processes rather than relying on a default assumption of isolation-by-distance (IBD), a pattern that is rarely validated and is often misunderstood in passively dispersed organisms across a wide range (76, 77). We provide a statistical approach that improves validation of these mechanistic relationships between environments and genomic patterns by assessing model performance and goodness fit relative to null expectations.

Though the LSR model has strong implications for the treatment of intraspecific genomic diversity, it is applicable beyond questions relating to single species. The apparent introduction of *G. granosissima* and persistence to date only in the Chesapeake Bay and Roanoke Sound indicate that LSRs may predict locations at which invasive species are more likely to persist, and boundaries past which they are less likely to move once introduced (33). These insights are generalizable to planktonic or passively dispersed species at large; they are cleanly concordant with simulated predictions of oceanographic effects on the distribution of genomic diversity across taxa (31, 35, 78–80). Oceanographic inference of LSRs should be considered as a primary mechanism for documented patterns of population structure in planktonic marine species.

The LSR model has the potential to explain the diversity and distribution of entire communities. Along the Atlantic Coast, species range boundaries cluster at Cape Hatteras, Cape Cod, and the Bay of Fundy, the boundaries predicted by LSR current modeling and captured by population genomic data in *G. demissa* (45). Clearly, a nonrandom assortment of boundaries has large ramifications for community diversity and is likely to be maintained by a contemporary mechanism. Predictions of range boundary locations are not concordant with sea surface temperature shifts, but do correspond with disruptions in ocean current flow, the kind that generate LSRs due to a dearth of propagules flowing in from upstream (45). The tendency for genetic patterns to co-localize with interruptions in connectivity is often noted (81, 82). Here, we validate ocean currents interacting with coastal complexity as a predictor for such patterns in coastal species, indicating generalizability from intraspecific genomic transitions to community distributional patterns that define an ecosystem.

When considered in full, this work indicates the need for new null expectations when examining diversity in the ocean and in passively dispersed organisms more generally: the creation and maintenance of LSRs, a source-sink dynamic (31, 34) where interruptions in dispersal create upstream regions whose native diversity dominates that of downstream regions until yet another interruption occurs. We provide a new framework for understanding distributional patterns in marine ecology, from genomes to communities, with ramifications for questions regarding coastal ecosystems that become more urgent in a rapidly changing environment (83).

## Materials and Methods

### Tissue collection and and preparation

The collection of specimens and tissues of *Geukensia* are as detailed in (52, 57; also see Supplemental Information Part A, Supplementary Table 1). Genomic DNA was isolated using a Qiagen DNEasy Blood and Tissue kit (QIAGEN, Hilden, Germany) and cleaned with a custom SpeedBead mixture (84, file S8) to remove low molecular weight fragments prior to applying the Adapterama RADseq protocol (48) using the enzymes *MspI, BamHI*, and *ClaI*. All libraries were pooled in equimolar ratios, size-selected using a Pippin Prep, and sequenced on an Illumina Novaseq X Plus PE 150 platform by Novogene Co. Inc. (Beijing, China).

### Data handling and population genomics

Sequence data were demultiplexed using process_radtags (85) and checked for correct restriction sites; following quality assessment all fragments were trimmed to 139bp. Reads were assembled *de novo* for presumed *G. demissa* individuals, removing several Atlantic coast individuals found to have *G. granosissima* mitotypes in (56). We used STACKS v2.66 (85) with parameters m=3, M=4, n=4, selected by iterating across parameter combinations and identifying the range of assembly parameter values at which additional polymorphic loci plateau while minimizing loci with excess heterozygosity. One random SNP from each assembled polymorphic locus was used for downstream analyses.

We filtered loci with SNPfiltR (86), removing loci missing >30% of genotypes and then individuals missing >25% of loci from the *G. demissa* dataset. Outlier loci were detected via PCAdapt, a technique that uses multiple linear regression to filter loci whose z-scores are inflated relative to an initial PCA of all individuals across the range (87). To determine whether outlier loci were LSR-specific, we also performed PCAdapt on subsets of individuals in sites with predicted ancestry from each LSR.

Our primary goal was identification of hierarchical structure relative to a null expectation of isolation-by-distance. The two patterns may, of course, be intertwined (22). We performed a Mantel test to detect isolation-by-distance, even though the asymmetrical directionality of planktonic dispersal renders a Mantel test, which assumes stationarity in both matrices, more likely to yield a false positive (53). We also ran separate Mantel tests within each LSR-driven region, except those that had very few sampling locations within them. As another measure of genetic differentiation across space, we calculated pairwise Weir-Cockham F_ST_ across all locations. To describe any hierarchical population structure present along the range, we performed DAPC using *adegenet* in R with 100 PCs and 5 DAs retained (88, 89) and sNMF via R package LEA (90) on the filtered *G. demissa-*only dataset without outliers.

### LSR modeling

We use the Mercator Ocean 1/12° (∼6km graticule) global physical model (91) to identify multigenerational sources for lineages that are initiated at high density along the Atlantic coastal domain. Simulated larvae are parameterized with typical pelagic larval duration (PLD) for *G. demissa*. These mussels free-spawn gametes during the summer (50), which form larvae without known vertical behavior, though the final larval stage is found near the sea surface (92). Larvae recruit to the mid-intertidal - often to the base of cordgrass (*Spartina*), mangrove, or other *Geukensia* - after about 2-3 weeks (50, 93).

For modeling purposes, the habitat is assumed to be the two 1/12th of a degree patches adjacent to the coast, and extends from the southern tip of Florida to the Gaspé Peninsula, with a recognized habitat gap across the Bay of Fundy from Rockland, Maine to Yarmouth, Nova Scotia where *G. demissa* are sparse or not present (94). Numerical models only resolve features over multiple grid-cells, so including the closest two grid-cells to the coast best represents the coastal boundary layer (95). Here we present results for simulations of PLD of 18 days, but the results are known to be stable over a much broader parameterization (29); see Supplementary Tables 7-9).

Each modeled habitat patch - in this case, the 2 patches closest to shore - was initially occupied by a single individual founding a unique lineage. Running the model with higher patch capacity does not change the outcome at this scale (33). The modeled circulation fields are used to calculate connectivity between habitat patches by releasing from each patch a Lagrangian, surface trapped, particle every half day in April, May and June of 2007 to 2024, for a total of 3276 particles (paths available at https://github.com/JamiePringle/EZfate). Each model generation, larval dispersal is modeled for each individual by releasing 16 propagules chosen randomly from the 3276 paths calculated. Of all released particles which reach a patch of habitat, one was chosen randomly to survive and propagate its lineage into the next generation. The model was run for 600 generations, at which time the ancestral location of each remaining lineage was noted as a potential lineage source location. Because the model is stochastic, the model run was repeated 10^4^ times, and all source regions were recorded. Contiguous source regions where the initial diversity is retained in at least 3 replicates are grouped as an LSR.

### Comparative oceanographic-genomic analysis

We performed redundancy analysis (RDA) in *vegan* (96) on the filtered *G. demissa-*only SNP dataset to quantitatively assess the strength of the relationship between LSR ancestry prediction and neutral genetic divergence. In order to compare this relationship with those predicted by other features of the environment, we ran three models, one for each of three different explanatory variables: latitude, mean annual sea surface temperature (SST; 98), and ancestry predictions for each sampling location resulting from the LSR model described above. Each dataset was reduced to mean allele frequencies per site coupled with a mean value of the predictor variable corresponding to each site to avoid pseudoreplication. Temperature values were bounded to the nearest 1/4° graticule, and LSR predictions for the three nearest neighbor habitat patches to the sample locations. Model performance was assessed via adjusted R-squared and pRDAs using function *anova.cca* in order to appropriately penalize the LSR model for including multiple distinct LSR ancestries as predictors. Given that predictors are highly correlated, backwards stepwise model selection was not feasible.

To generate a genomic dataset representing the neutral expectation of isolation by distance for comparison to our genomic data, we used SLiM (99) to simulate genotypes in a spatially explicit, stepping-stone isolation by distance model. Given a spatially explicit model of IBD, simulations in SLiM were tuned via a migration parameter to generate either a range-wide F_ST_ of 0.03 (the divergence of all the unglaciated range, see Results), 0.10 (the divergence across the entire range), or a midpoint value (see Supplementary Figure 19). Simulated 6Mbp chromosomes were sampled following 10^4^ generations of evolution under stepping-stone migration, mutation rate of 10^-8^, population sizes of 1000, and all mutations having no fitness effect. One hundred replicate simulations were generated for each migration parameter, and the resultant VCF files for each replicate were converted to .geno format (90) and assessed using sNMF with the same value of K as our empirical data.

We use probability distributions for ancestry from our empirical genomic sampling, from the LSR model, and the SLiM IBD simulations to calculate the goodness of fit of empirical data to the LSR model and to IBD. LSRs that contributed less than 5% to the predicted ancestry of our sampled locations were ignored, and when there remained more LSR vectors than the value of K for genomic data, LSR predictions with high spatial overlap were combined. Probability vectors in all 3 data types were re-organized from south to north using the centroid of their predicted range so that all were directly comparable in space. The distance from observed genomic ancestries to the LSR model or IBD simulations were measured using Hellinger distances at each location, where the root mean squared error was calculated across locations to identify the predictive fit of IBD or LSR models because locations are not statistically independent from one another.

## Author Contributions

CAS performed research, analyzed data, and wrote the paper; KNP performed research, analyzed data, and wrote the paper; JMP performed research, contributed new analytic tools, analyzed data, and wrote the paper; JPW designed research, performed research, analyzed data, and wrote the paper.

## Acknowledgments

We appreciate the resources and people associated with the “Evolving Seas” Research Coordination Network supported by the National Science Foundation. We benefited from collecting assistance from David Gaspard, Tanya O’Reilly, Katie Miller, David S. Johnson, Roland Hagan, Thomas Grothues, Jenny Shinn, April Blakeslee, John Carroll, Jim Morley, Juliet Wong, Lauren Baena, Ken Halanych, Riley Wares, and the Marine Biological Laboratory. Funding support to JPW from the UGA Vice President for Research, Elaine Lutz Foundation for Aquatic Biodiversity, Katelyn Chandler Memorial Award, and the Odum School of Ecology. Funding support to JMP from National Science Foundation OCE-1947954. Additional technical and writing guidance from Morgan Kelly, Katie Lotterhos, Seth Wenger, and Erik Sotka was greatly appreciated.

## Supplemental Information

Supplemental results and data are presented below.

## Supplemental Information

### Part A. Sampled locations

**Supplementary Table 1:**
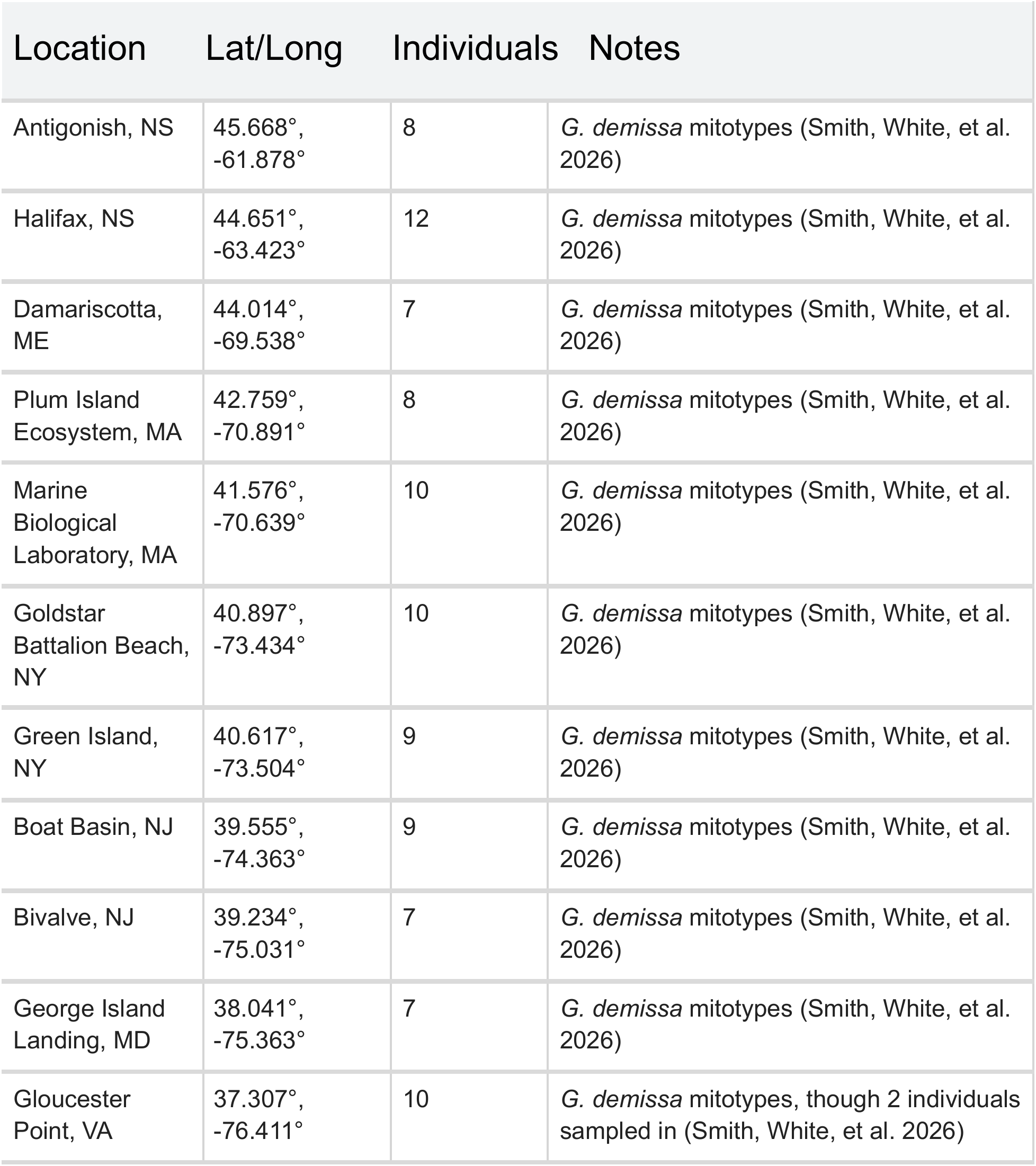
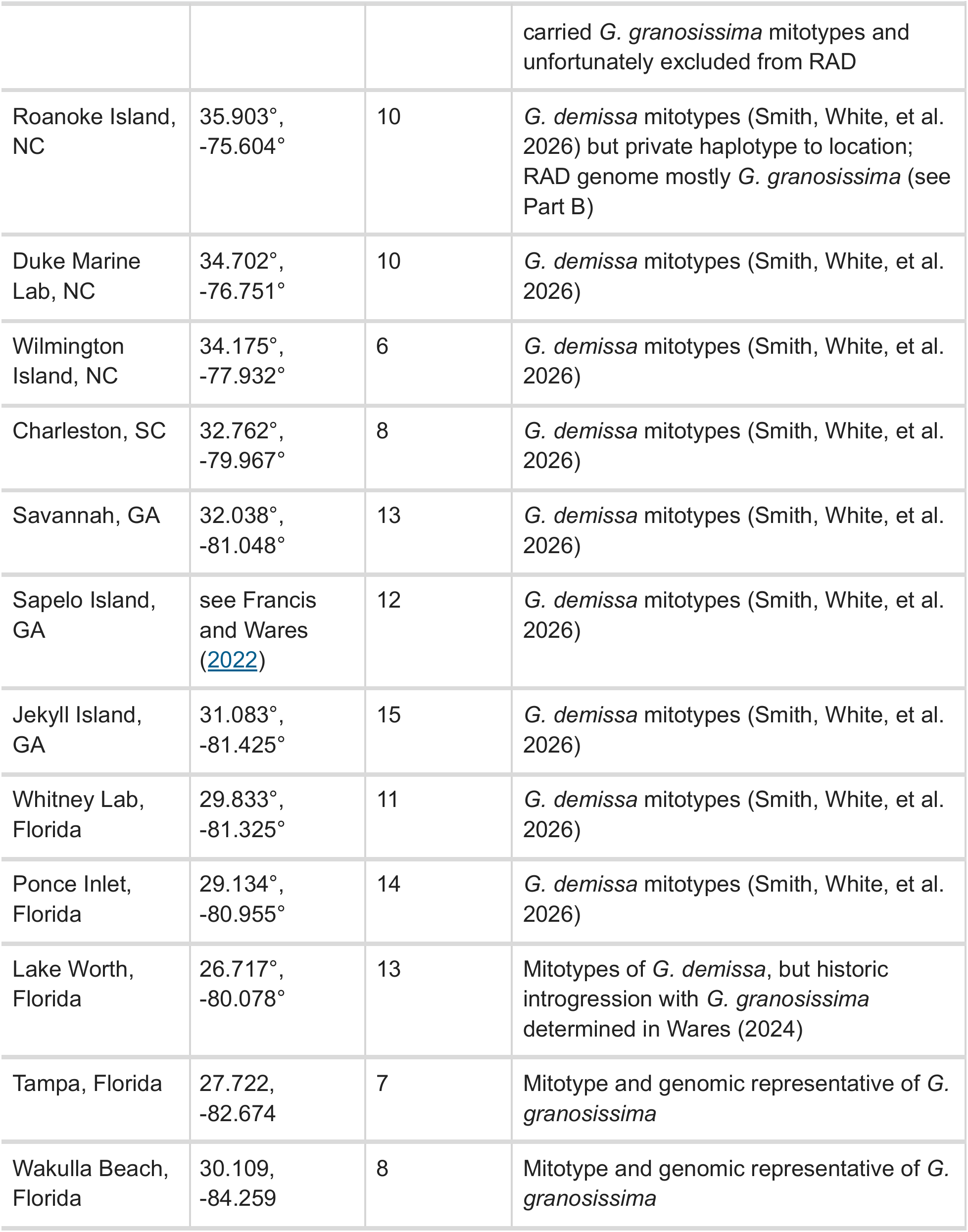
Sampling locations of *G. demissa* and *G. demissa* samples.

### Part B. Examining data from the focal *Geukensia demissa* and the Gulf congener *G. granosissima*

The complete genomic data set for this study includes sequenced RAD libraries from Wares (2024) as well as new sequence data. The combined spatial breadth involves samples from Wakulla Beach, FL in the Gulf of Mexico to Antigonish, Nova Scotia, Canada. Collections from the Gulf of Mexico carry the mitotype and distinct genome of *G. granosissima* (1, 2). A small amount of *G. granosissima* diversity had been found in southeastern Florida; the Gulf species was historically distributed on the Atlantic coast as well but little trace remained, per (1).

All libraries were prepared using the Adapterama RAD protocol of (3) in the Wares lab at the University of Georgia. All pooled libraries were sequenced by Novogene in batches submitted in 2023 (1), 2024, and 2025. Data are available from the Short Read Archive at NCBI accessions PRJNA977713. All individuals sequenced have COI mitotypes archived at NCBI (4). Average read number per individual was 7.157 million reads across 224 individuals; the Atlantic-only data assembled and filtered as a single species comprised 6.951 million reads across 195 individuals.

The sequence data including both species were tested for optimal parameterization of assembly in Stacks 2 (5) using a custom workflow developed by Rachel Toczydlowski and colleagues (used with permission). The parameters chosen for assembly of the complete dataset were M=6, n=7 as the two mussel taxa in our data are approximately 4my divergent, limiting the number of loci that share restriction sites and homology (6). Both the all-sites and random-snp VCF outputs were selected from the populations module in Stacks.

### SNP Filtering

The VCF data were imported into RStudio and filtered according to two schemes, given a large amount of missing data in the matrix (again, in part because many variable loci are only found in one species or the other; see plot below). Out of 224 individuals sequenced, only about 20 were known *a priori* to be *G. granosissima* (samples from Tampa Bay, Florida and Wakulla Bay, Florida). These individuals had canonical *G. granosissima* mitochondrial haplotypes as well as a high probability of genotype background distinct from that of Atlantic coast individuals (1). **Filtering approach A** removed any loci that retained less than 75% of genotype calls (using the R package SNPfiltR (7), and then any individual missing more than 95% of genotype calls. This retained 4053 SNPs out of 715,438 in the original data file and only eliminated 3 individuals. **Filtering approach B** filtered out loci retaining less than 30% of genotypes, leaving 32,718. We then filtered by sample (0.95) and again by SNP (0.6) to retain 11,811 loci.

**Supplementary Figure 1:**
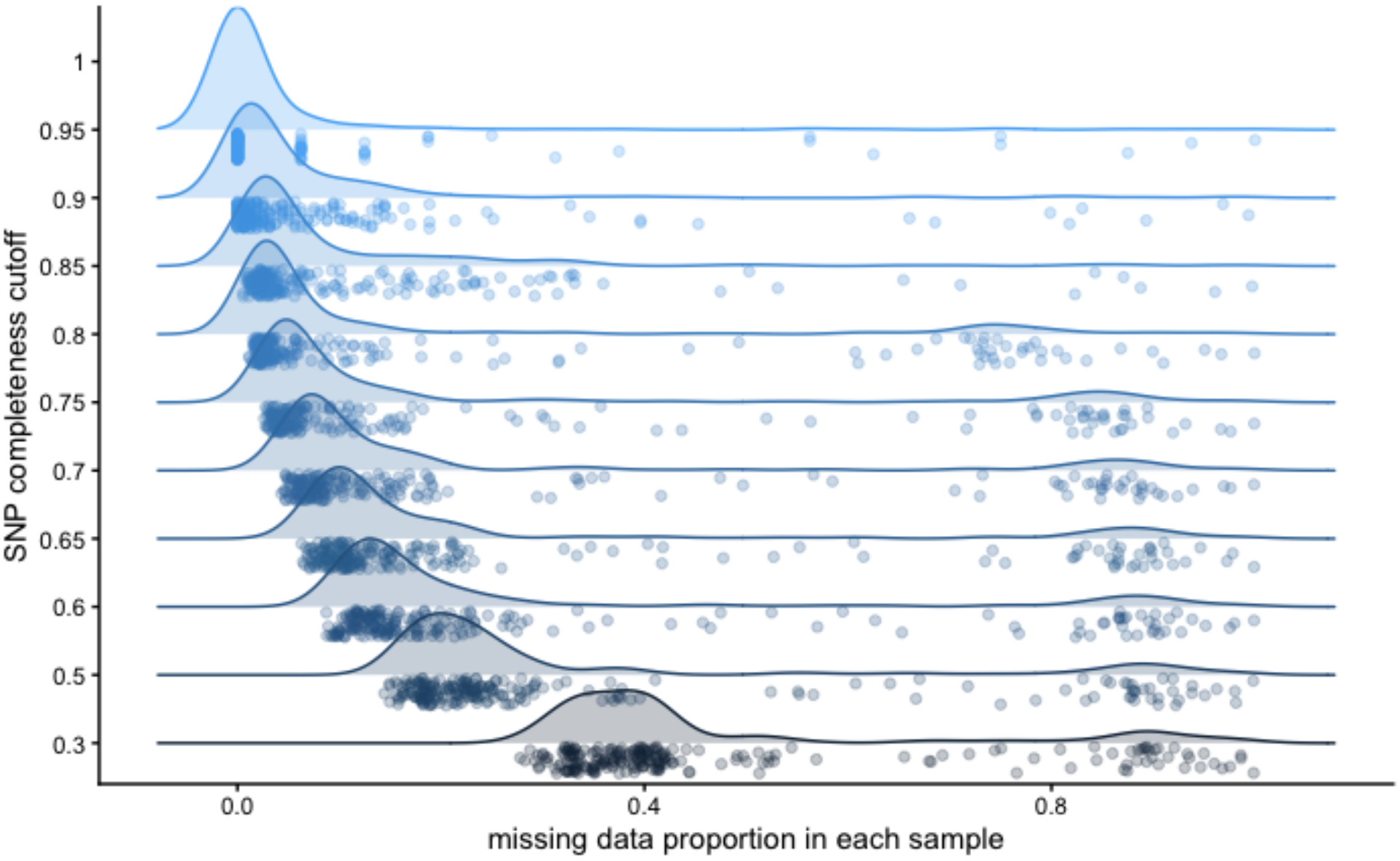
Density plot of the proportion of SNPs missing in each sample after applying varied SNP completeness cutoffs during the filtering process. *Note the two clusters of “missingness” correspond to the two species included in analysis*.

VCFs were converted to genind format for analysis using the package adegenet (8). We first evaluated for batch effects using a Discriminant Analysis of Principal Components (9); however our batches are largely (but not entirely) non-overlapping geographically. We do see concordance between 2024 and 2025 spatial samples from the Canadian Maritimes, and 2025 sequencing of Roanoke Sound individuals align with 2023 sequencing of Florida populations of *G. granosissima* - a first sign of an unusual element of our results.

**Supplementary Figure 2:**
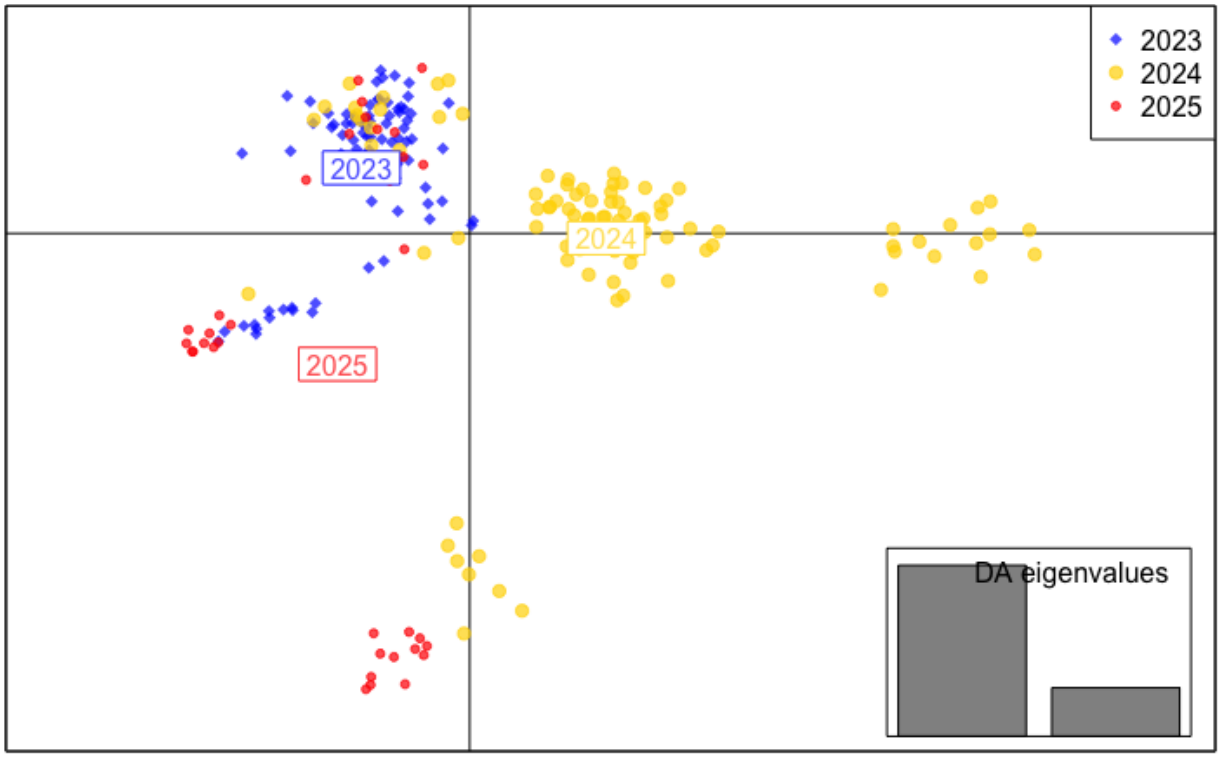
A DAPC plot of all *G. demissa* and *G. granossissima* samples, with point color representing the year in which they were sampled.

Both filtering approaches were subsequently evaluated with DAPC (9) retaining 3 principal components and using 100 discriminant functions each. We note that Filtering Approach A (fewer loci retained) clearly separates samples from the Canadian Maritimes and Gulf of Maine on principal components 1 and 2, but does not clearly separate the Gulf of Mexico species from the Atlantic. The Gulf of Mexico and Atlantic species are instead separated between PC 1 and PC 3 (as is the Roanoke sample).

**Supplementary Figure 3:**
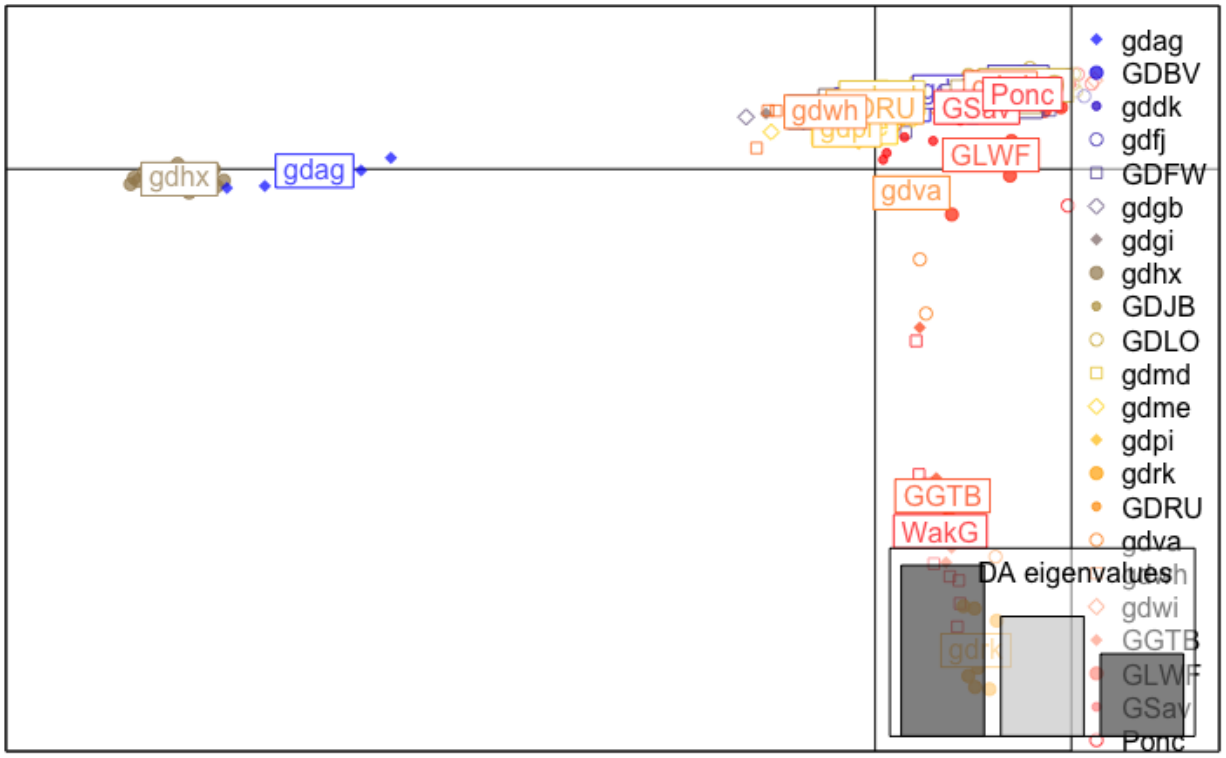
A DAPC plot of samples from *G. demissa* and *G. granossissima* under filtering approach A. Colors represent sampling locations from northern latitudes (blue) to southern latitudes (red).

Our second Filtering Approach B more clearly isolates *G. granosissima* and the Canadian samples from the rest of *G. demissa* on PC 1 and PC 2, with the Gulf of Maine separating out on PC 3.

**Supplementary Figure 4:**
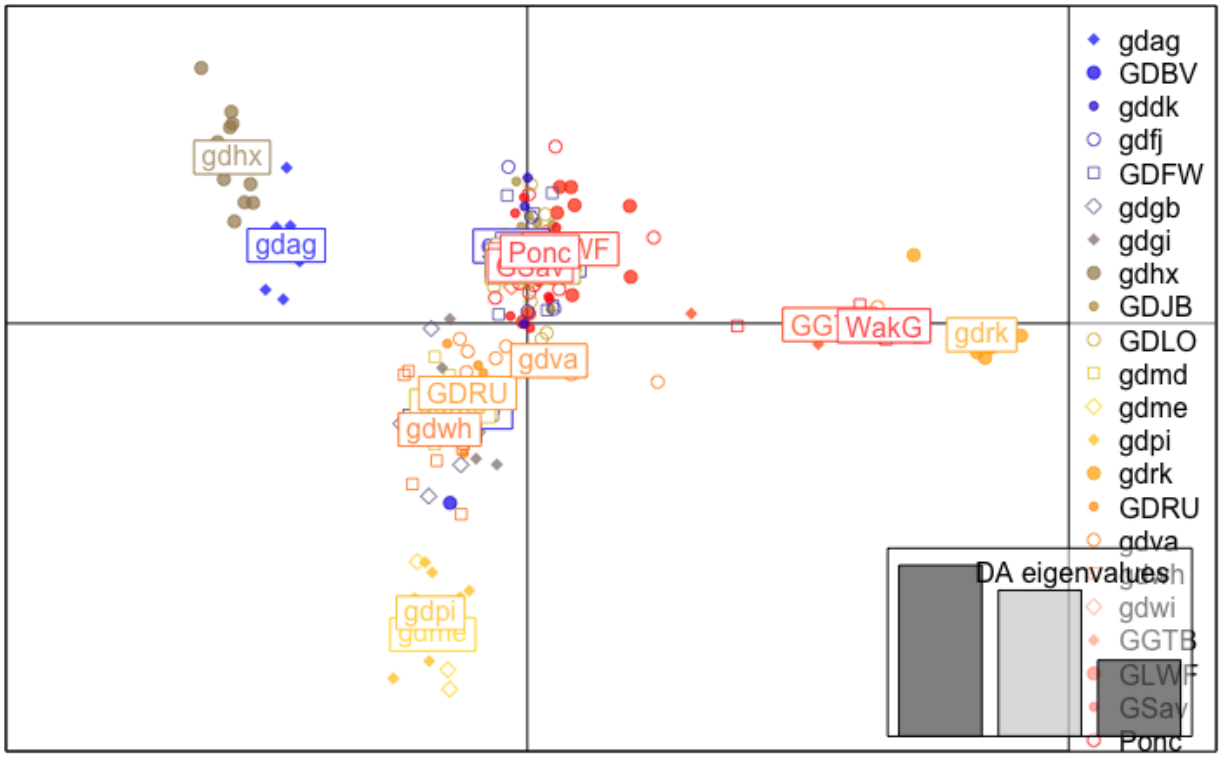
A DAPC plot of samples from *G. demissa* and *G. granossissima* under filtering approach B. Colors represent sampling locations from northern latitudes (blue) to southern latitudes (red).

Following this, data from both filtering approaches were analyzed using snmf to explore population structure given K ancestral lineages. In each case, 20 repetitions for K=1 to 10 were explored for cross-entropy to help select the most appropriate value(s) of K for visualization. Filtering Approach A suggests that K=4 is the objectively highest number of ancestral lineages given the cross-entropy of the data.

**Supplementary Figure 5:**
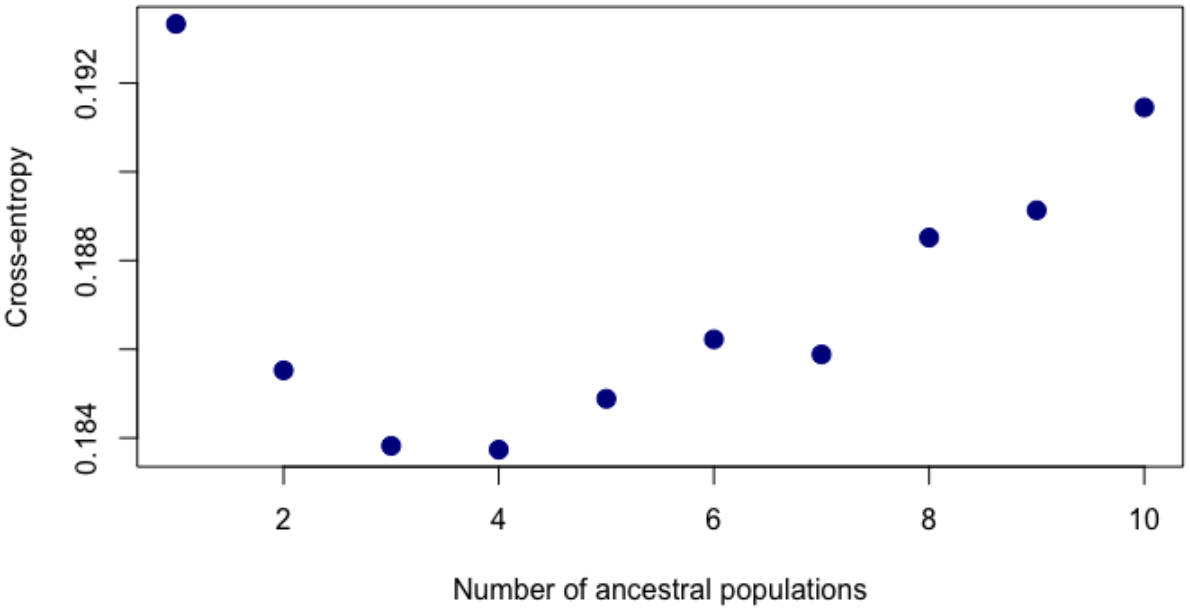
Cross-entropy plot of the dataset including *G. demissa* and *G. granosissima* under filtering approach A as a function of *k* ancestral populations. Lower cross-entropy represents more stable matrix factorizations.

The snmf results for K=2 through 5 with filtering approach A are presented at right. The key observation is that the samples at furthest left represent Wakulla and Tampa in the Gulf of Mexico, and match likely ancestry with samples from Roanoke Sound, NC and (partially) 2 samples from Virginia. The right-most cluster of individuals are from the Canadian Maritimes, preceded by the Gulf of Maine. All samples are presented in latitudinal “order” starting in the Gulf of Mexico and moving towards Canada as you proceed to the right. Colors are not consistent among runs. “Missingness” of genotypes is plotted at the bottom of this panel.

For Filtering Approach B (more loci), K=3 or 4 is again the most objectively optimal outcome. Results for K=2 through 5 are presented. Based on these results, we continued analysis of the data only from the Atlantic coast of North America as representing *G. demissa*, but with the samples from Roanoke Sound representing apparently introduced diversity (see main manuscript).

As a final exploration of the effect of ‘missingness’ on this pattern, we filtered a third way; most individuals are missing more than 90% of the >715k SNPs in the STACKS assembly; filtering at this level leaves only 159 SNPs but more even missingness across the two species. With this final approach, cross-entropy testing suggests the best value of K for so few data is 2; higher levels lead to obvious over-fitting of data, and the answer is unchanged in terms of the apparent introduction of *G. granosissima* to the Chesapeake Bay and Roanoke Sound.

In addition to these clustering analyses, the pairwise Fst values among sites in this full data set also indicate high similarity between the Roanoke sample and our Gulf of Mexico samples in (1). Pairwise Fst among most Atlantic samples and the Gulf populations averages 0.515, while Wakulla to Tampa is 0.024 and either of those samples to Roanoke is 0.113. Values of Roanoke to other Atlantic coastal samples are uniformly high (0.381).

**Supplementary Figure 6:**
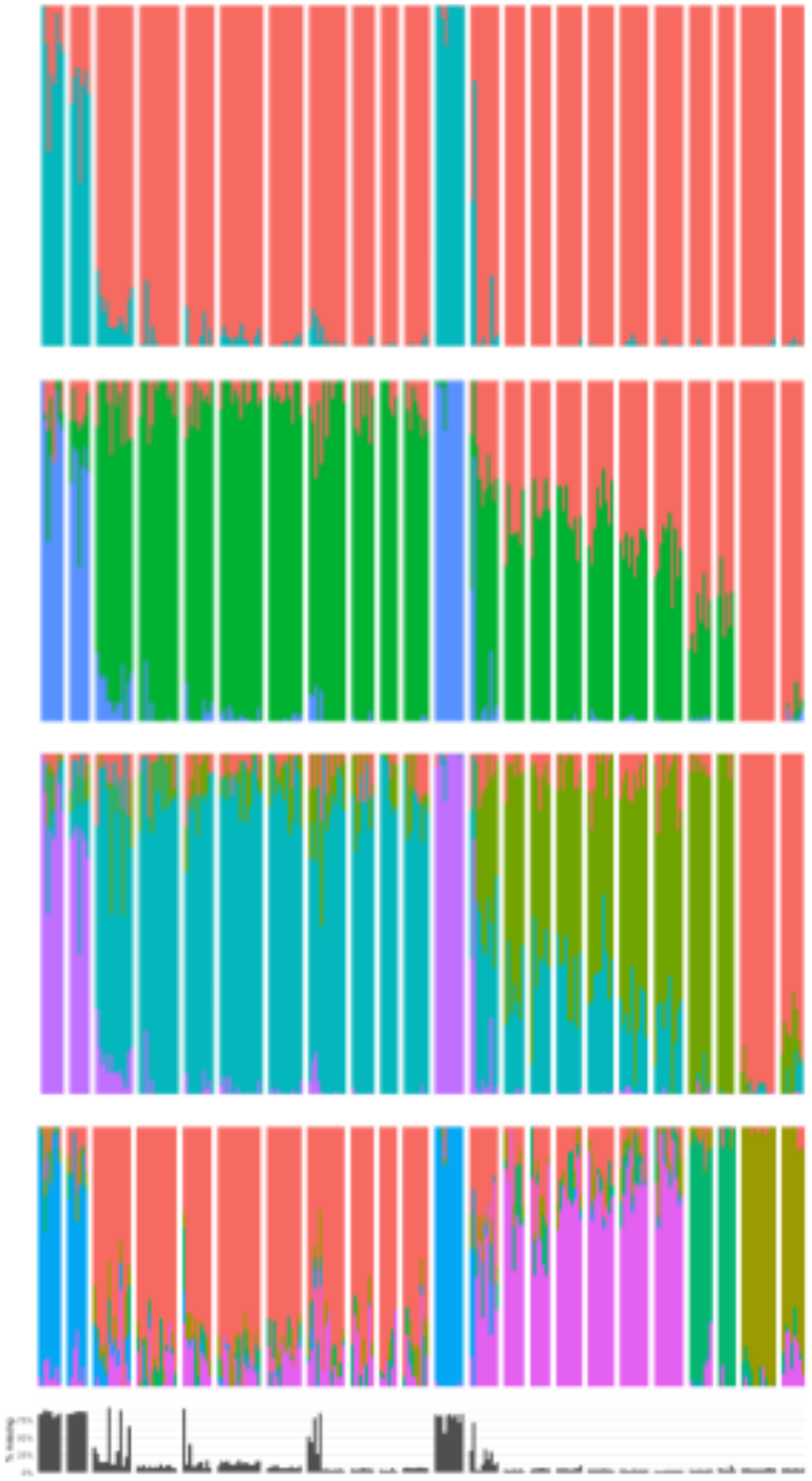
SNMF plots of the *G. demissa* and *G. granosissima* dataset under filtering approach A with different values of *k* ancestral populations of that individual’s SNPs derived from each ancestry group. Gray bar plot represents proportion of missing SNP data in each individual.

Overall this appears to be a subset of Atlantic samples (all of Roanoke, some of Gloucester Point) that - following DAPC and snmf - appear to be anthropogenic introduction of *G. granosissima* to this region followed by hybridization and biased introgression towards establishment of a regional population of *G. granosissima* with mitochondrial capture of an endemic *G. demissa* mitotype. This type of discordant introgression has been seen in other species introduction events (10), but further sampling of this region will be needed to dissect the dynamics. Calculation of nucleotide diversity (π) from our data – though a biased measure given this is RAD data and not an all-sites VCF (see Section C) – suggests much lower diversity at Roanoke (π of 0.032) relative to the rest of the Atlantic coast (π 0.0632). F is is much higher at Roanoke (0.452) than at other Atlantic coastal sites (mean 0.242). Additionally, estimates of linkage disequilibrium (*r_d_*) are elevated at Roanoke and Virginia sites relative to surrounding areas, and Hamming distances among individuals are far lower at Roanoke.

**Supplementary Figure 7:**
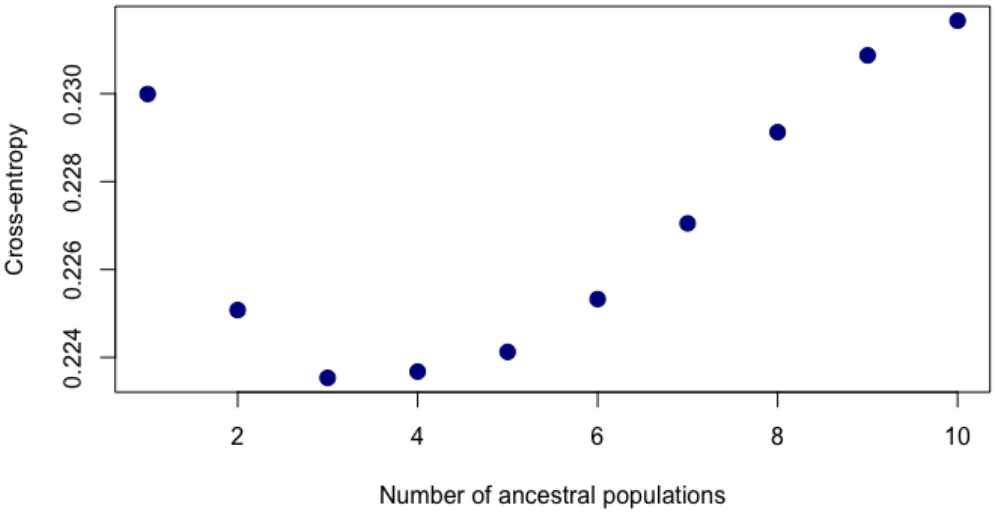
Cross-entropy plot of the dataset including *G. demissa* and *G. granosissima* under filtering approach B as a function of *k* ancestral populations. Lower cross-entropy represents more stable matrix factorizations.

The source of this introduced diversity is not known, but there is evidence of oysters (*Crassostrea virginica*) from Louisiana being transported to the Chesapeake region in recent decades (11–13). Indeed there are other examples of movement of Louisiana oysters to the North Carolina coast (14).

**Supplementary Figure 8:**
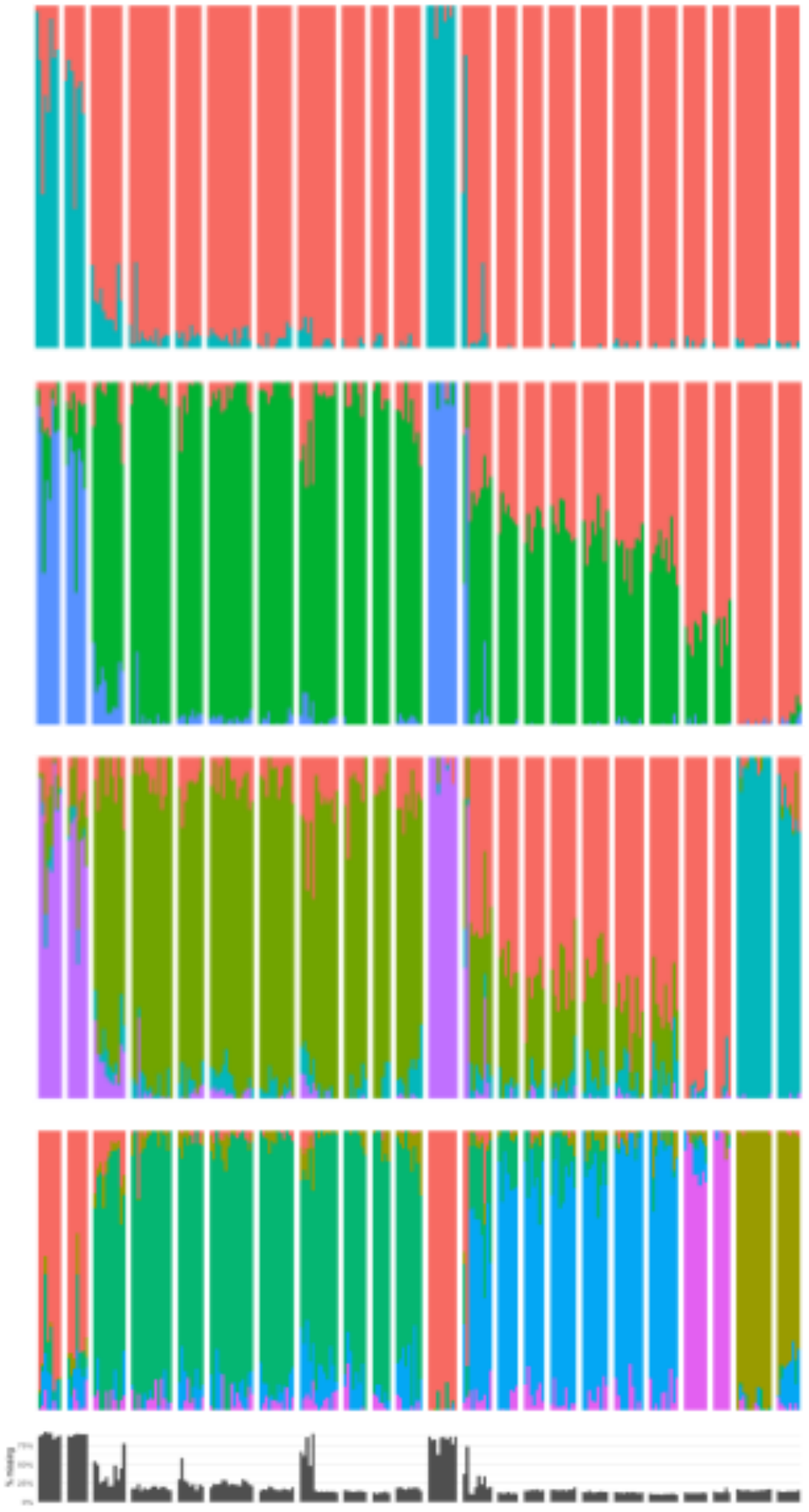
SNMF plots of the *G. demissa* and *G. granosissima* dataset under filtering approach B with different values of *k* ancestral populations. Bars represent individuals, and bar color represents the proportion of that individual’s SNPs derived from each ancestry group. Gray bar plot represents proportion of missing SNP data in each individual.

**Supplementary Figure 9.**
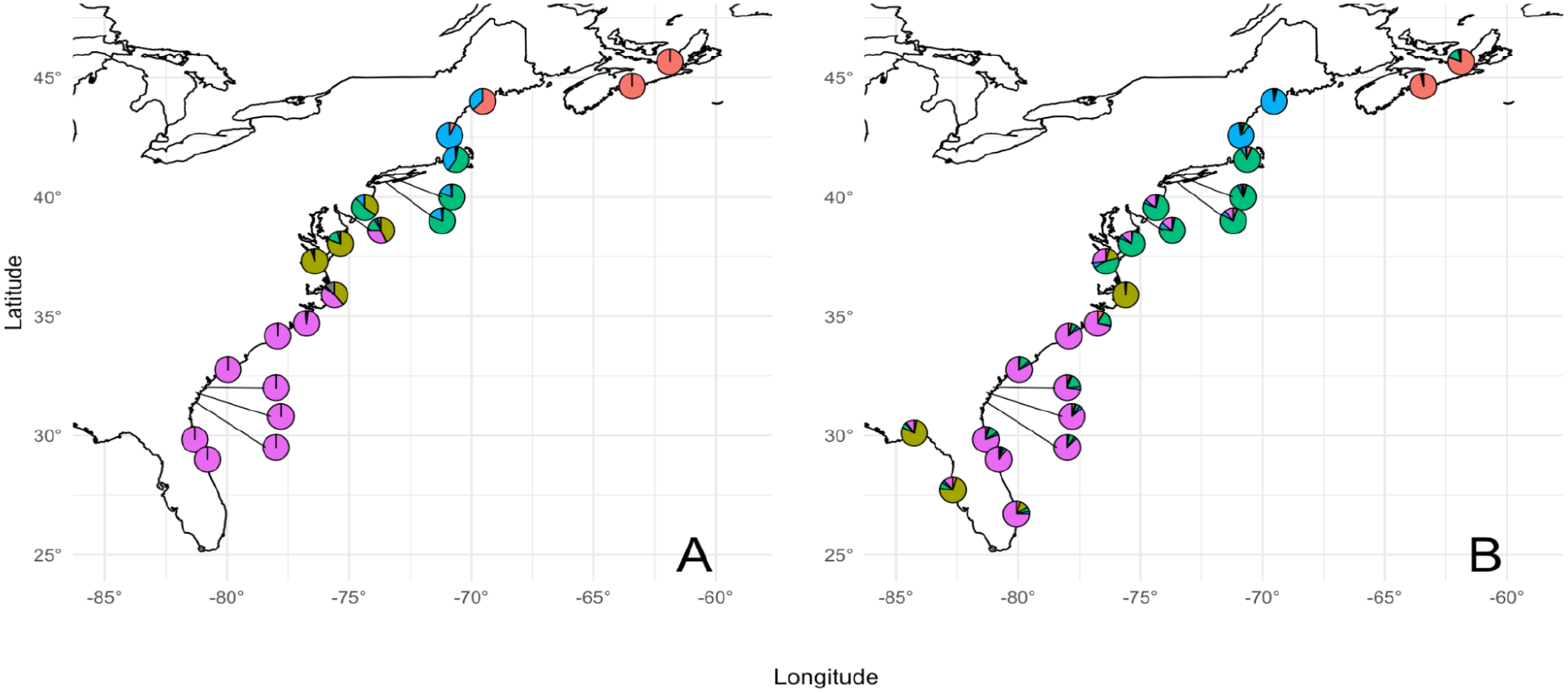
Map of sampling locations in which pie chart colors represent predicted genomic ancestry via ocean current modeling (A) and mean snmf-assigned ancestry group for filtering regime B in the *G. granosissima*-inclusive dataset (B). Though the optimal number of clusters *k* is 4, *k = 5* is depicted to display *G. granosissima* introgression.

### Part C. Mitochondrial diversity

COI mitotypes are available at Genbank for all individuals in this study (4). A minimum spanning tree from (4) is replicated below showing the relationship of the private haplotype for all Roanoke individuals relative to all other sequences from *Geukensia demissa*. As noted in (4), we have also detected 5 individuals on the Atlantic coast that carry a *G. granosissima* mitotype (which is ∼ 12% divergent from the *G. demissa* sequences and is found in all individuals sampled along the Gulf of Mexico coast to date): one individual from Ponce Inlet (1), 2 individuals from our Gloucester Point site, and 2 individuals from Roanoke Sound site. These latter individuals were not prepared for RAD sequencing; we did not know a mechanism for those mitotypes to be in that region at the time. Chromatograph data are available in CodonCode Aligner format or raw .ab1 upon request from JPW; the single diagnostic site supporting the private mitotype at Roanoke is high quality (Phred > 30) in all individuals. Note that in this network haplotypes are pooled into 1, but there are actually multiple private mitotypes in the Roanoke population.

**Supplementary Figure 10:**
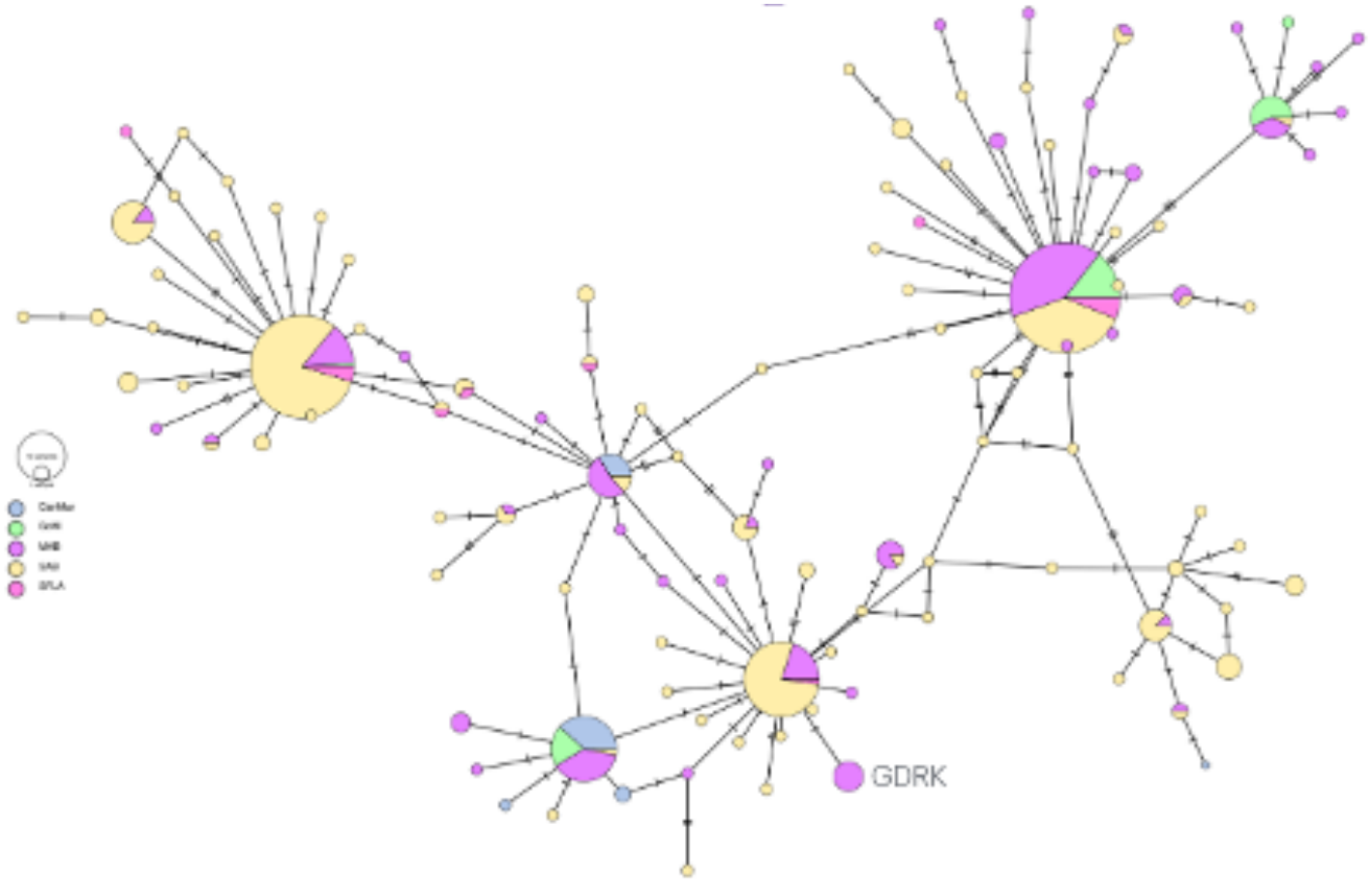
Haplotype network representing mitochondrial DNA from *G. demissa* samples collected in (4). Pie color represents the proportion of a haplotype that arises from each major marine ecoregion (light blue for the Canadian Maritimes, light green for the Gulf of Marine, purple for the Mid-Atlantic Bight, yellow for the South Atlantic Bight, and pink for Southern Florida.

Additionally, we present a neighbor-joining tree of all available mitotypes from our studies (1, 4) showing the divergence of *G. granosissima* mitotypes in dark red (all sequences are from the Gulf except 1 from Ponce Inlet, 2 from VA, and 2 from Roanoke) and the private haplotype of Roanoke Sound (pink).

**Supplementary Figure 11:**
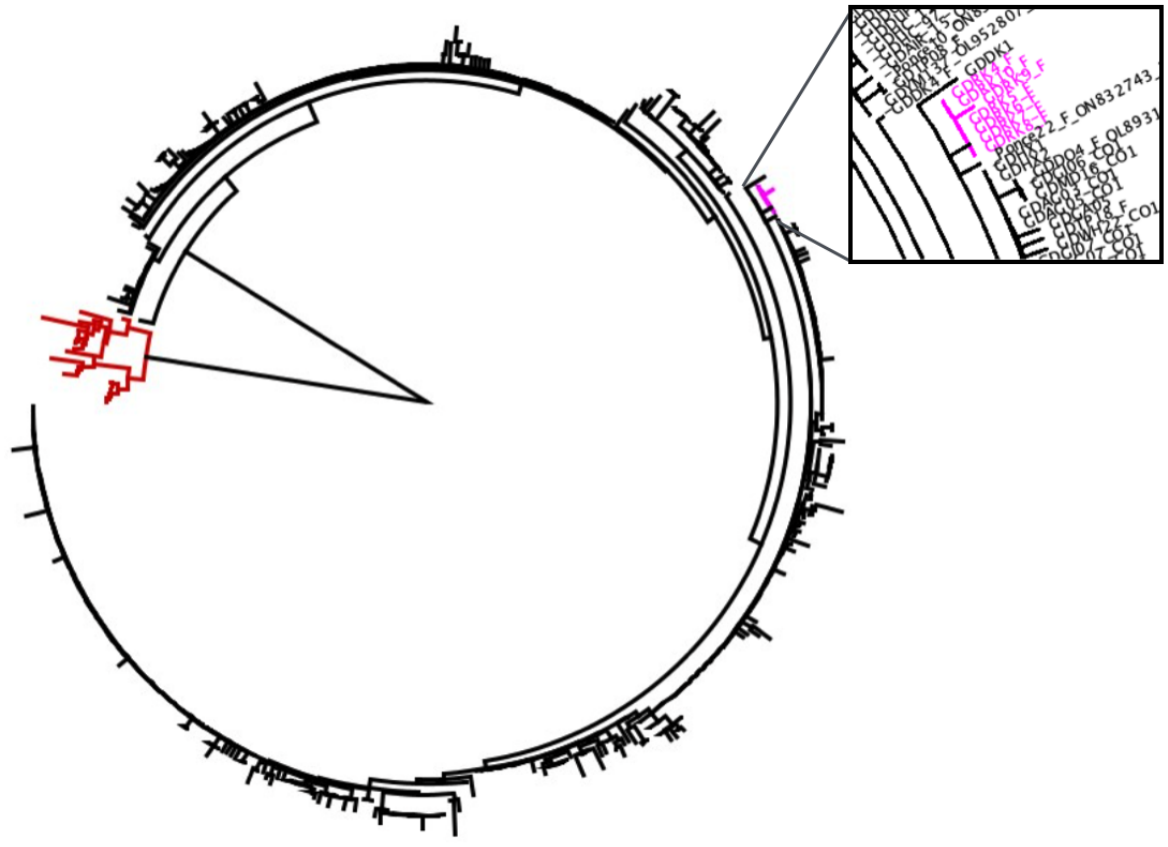
Neighbor-joining tree representing mitochondrial samples collected in (1, 4). The private haplotype from Roanoke Sound is represented in pink, and samples from the Canadian Maritimes are colored red.

### Part D: Outlier screening and analysis

PCAdapt (15) was performed on G. demissa and G. granosissima datasets with an a-level of 0.05. All putative outliers were excluded.

**Supplementary Figure 12:**
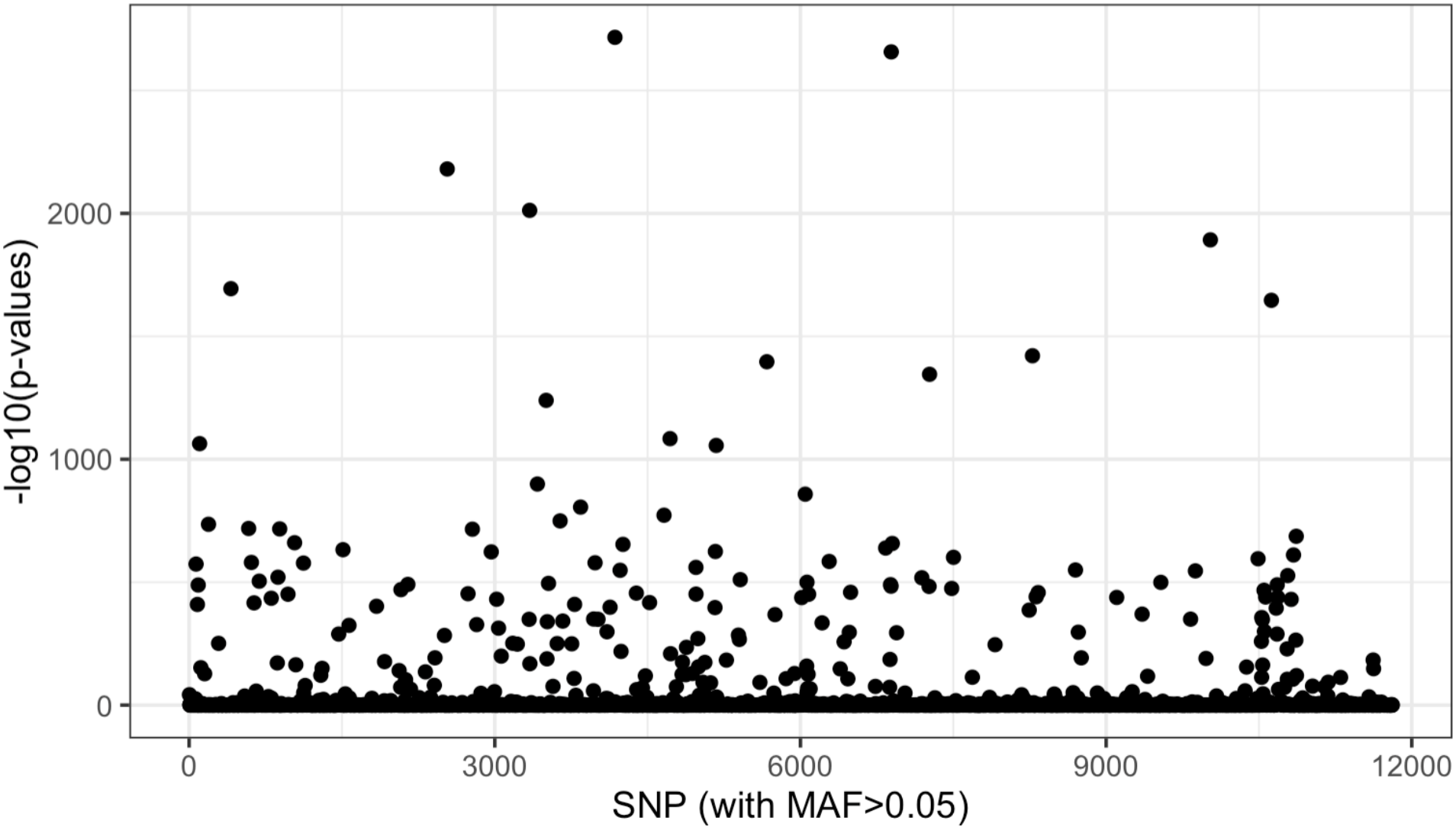
A Manhattan plot of outlier loci for the *G. demissa* and *G. granosissima-*inclusive dataset. High values on the y-axis indicate that allele frequencies at a given locus are unlikely to be explained by neutral population structure. High numbers of outliers as seen on the plot indicate high missingness in data, a common occurrence when reads from two distinct species are assembled in the same STACKS run. In total, PCAdapt detected 399 outliers in this dataset, indicating both large swaths of missing data and divergence between *G. demissa* and *G. granosissima* individuals.

**Supplementary Figure 13:**
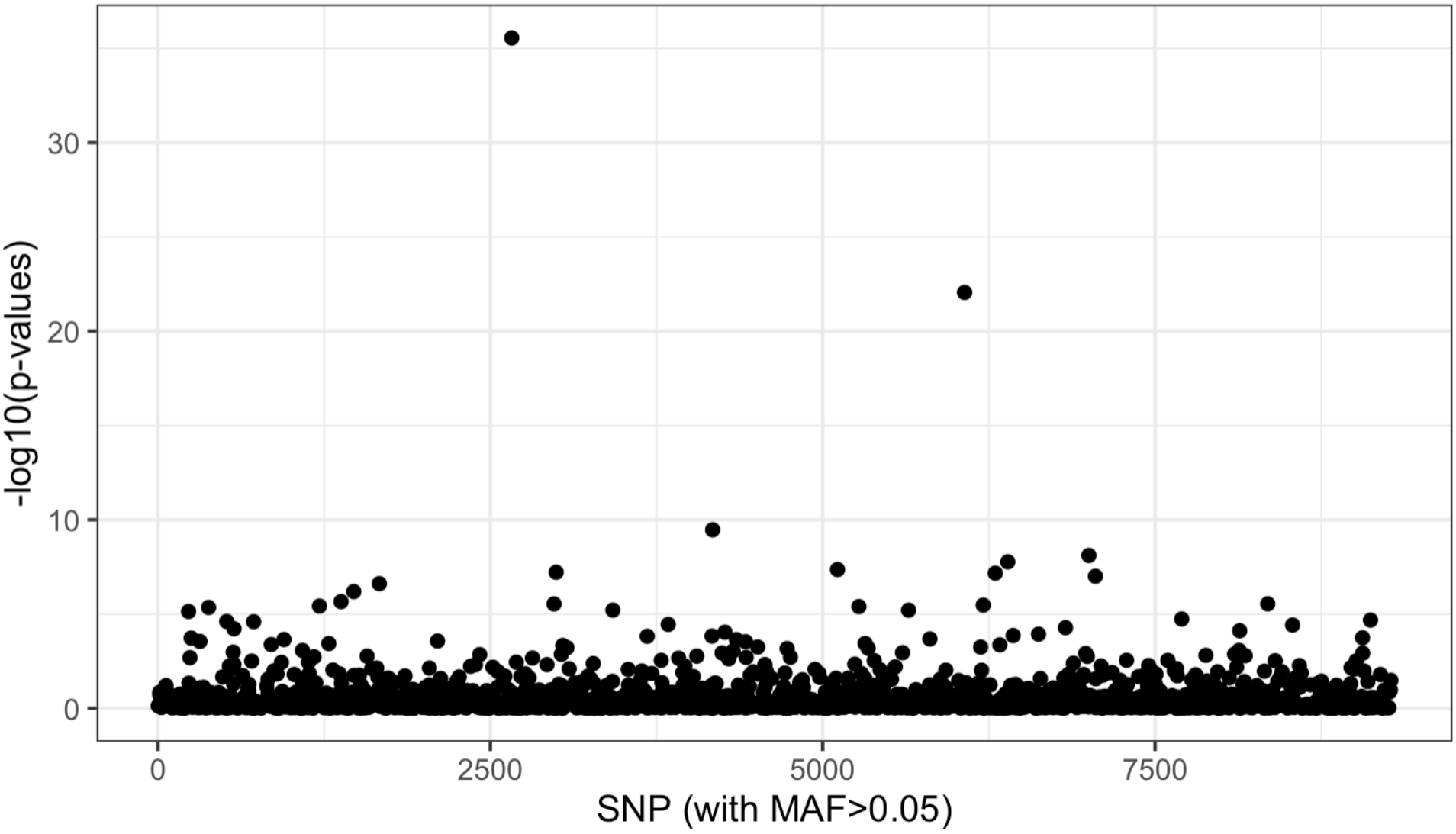
A Manhattan plot of PCAdapt p-values for the entire *G. demissa* dataset. High values on the y-axis indicate that allele frequencies at a given locus are unlikely to be explained by neutral population structure.

**Supplementary Table 2: Outlier loci lists for *G. demissa* [Gdem_outliers.txt](https://drive.google.com/file/d/1ugx3u8pdYDe7mGpMYOm51A1WovQS4JCo/view?usp=sharing)**

**Supplementary Table 3: Outlier loci for the Gulf of Maine [Maine_outliers.csv](https://drive.google.com/file/d/177WJwGDixQ4je4hMTJ2lgdqS3zCmYu_P/view?usp=sharing)**

**Supplementary Table 4: Outlier loci for the Mid-Atlantic Bight [Mab_outliers.csv](https://drive.google.com/file/d/1Kq6D5-8gkQMtcbkMwsD3hDVRQKUvMzmn/view?usp=sharing)**

**Supplementary Table 5: Outlier loci for the South-Atlantic Bight [Sab_outliers.csv](https://drive.google.com/file/d/1Upl-rjaO1jXon4VkGla2JFFfnjoEO_od/view?usp=sharing)**

**Supplementary Table 6: Outlier loci for the Canadian Maritimes [maritimes_outliers.csv](https://drive.google.com/file/d/1XUCtDYiNt28Xo_ayiQbj5T9PJ5dCPSaZ/view?usp=sharing)**

### Part E: DAPC and sNMF

DAPC identified and retained 5 clusters for the *G. demissa* dataset by comparing the Bayesian Information Criterion (BIC) of successive k-means clustering of the PCA-transformed dataset (9). DAPC is preferable in this dataset rather than PCA due to the high level of background variation expected in a hyperdiverse organism such as *Geukensia demissa* (1). Whereas PCA might be unable to identify clusters in the presence of such high expected variation, DAPC seeks to determine where variation occurs between subsamples, minimizing variation within assigned clusters and maximizing variation between them (9)

**Supplementary Figure 14.**
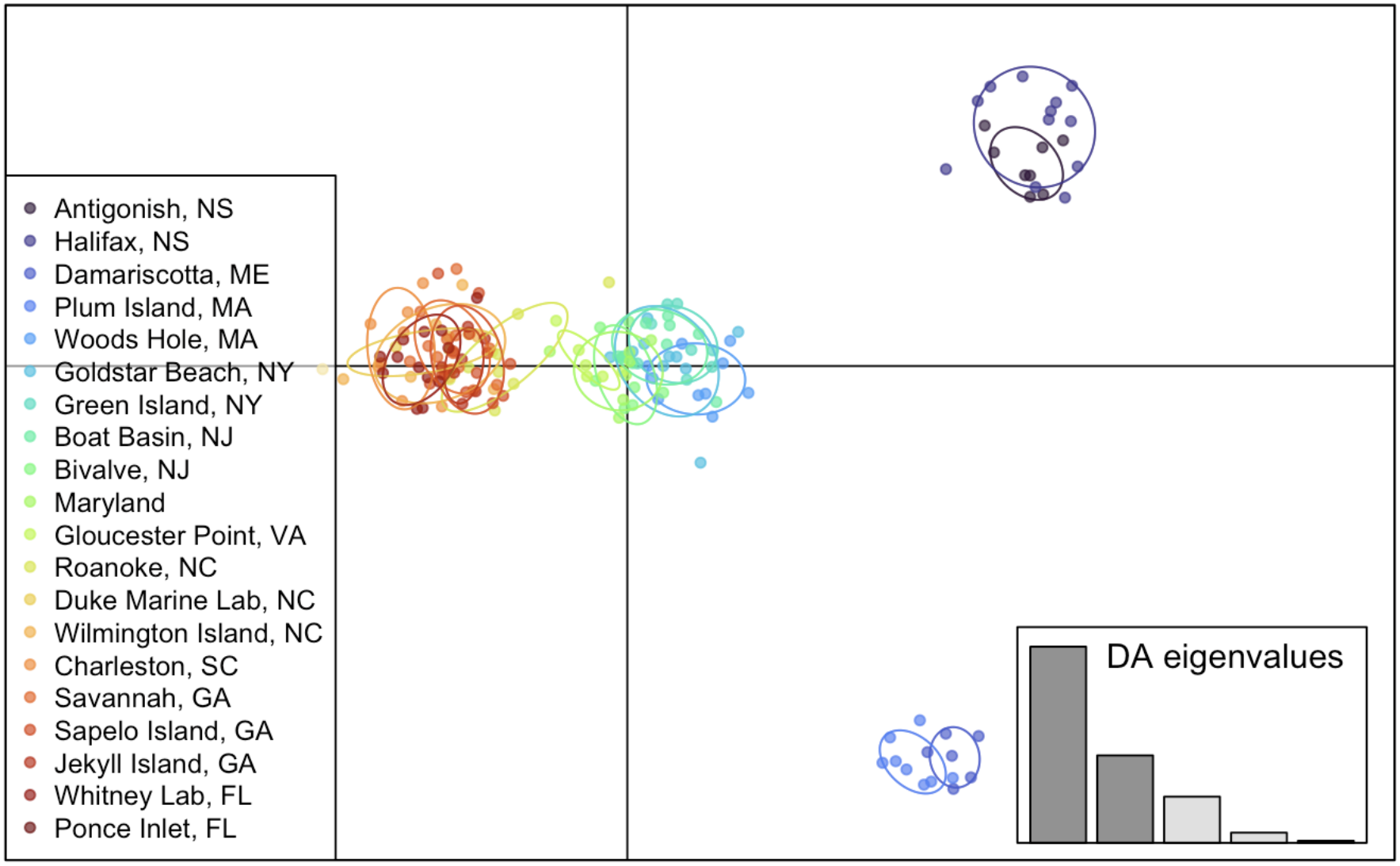
DAPC plot from the *G. demissa-*only neutral dataset, with cooler colors representing northern latitudes and warmer latitudes representing southern latitudes.The optimal number of clusters to retain was 5, with retained PCA axes capturing 7.98% of total variation.

**Supplementary Figure 15.**
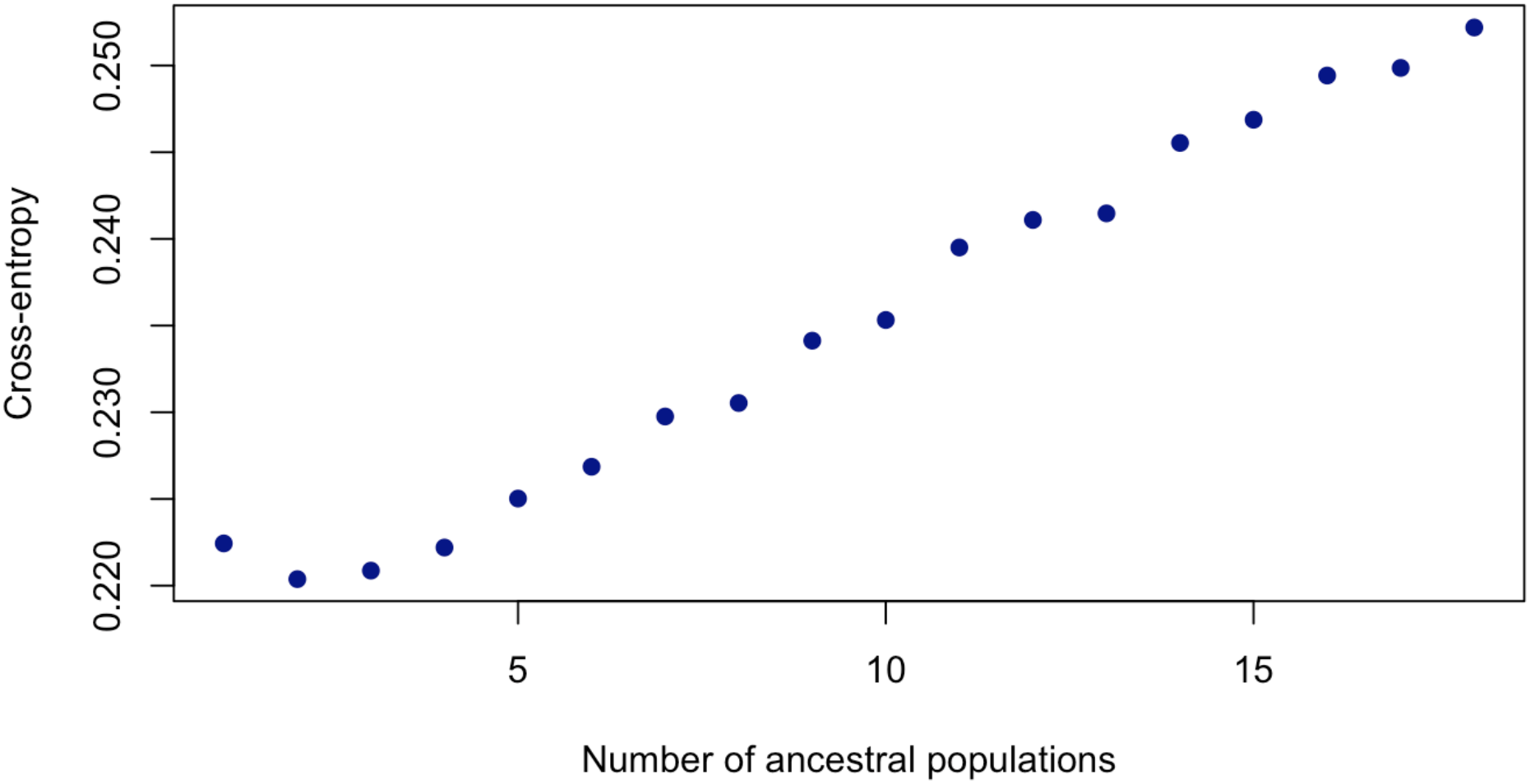
Cross-entropy plot of values of *k* ancestral populations for the *G. demissa* snmf run. Lower cross-entropies represent more stable matrix factorizations.

**Supplementary Figure 16.**
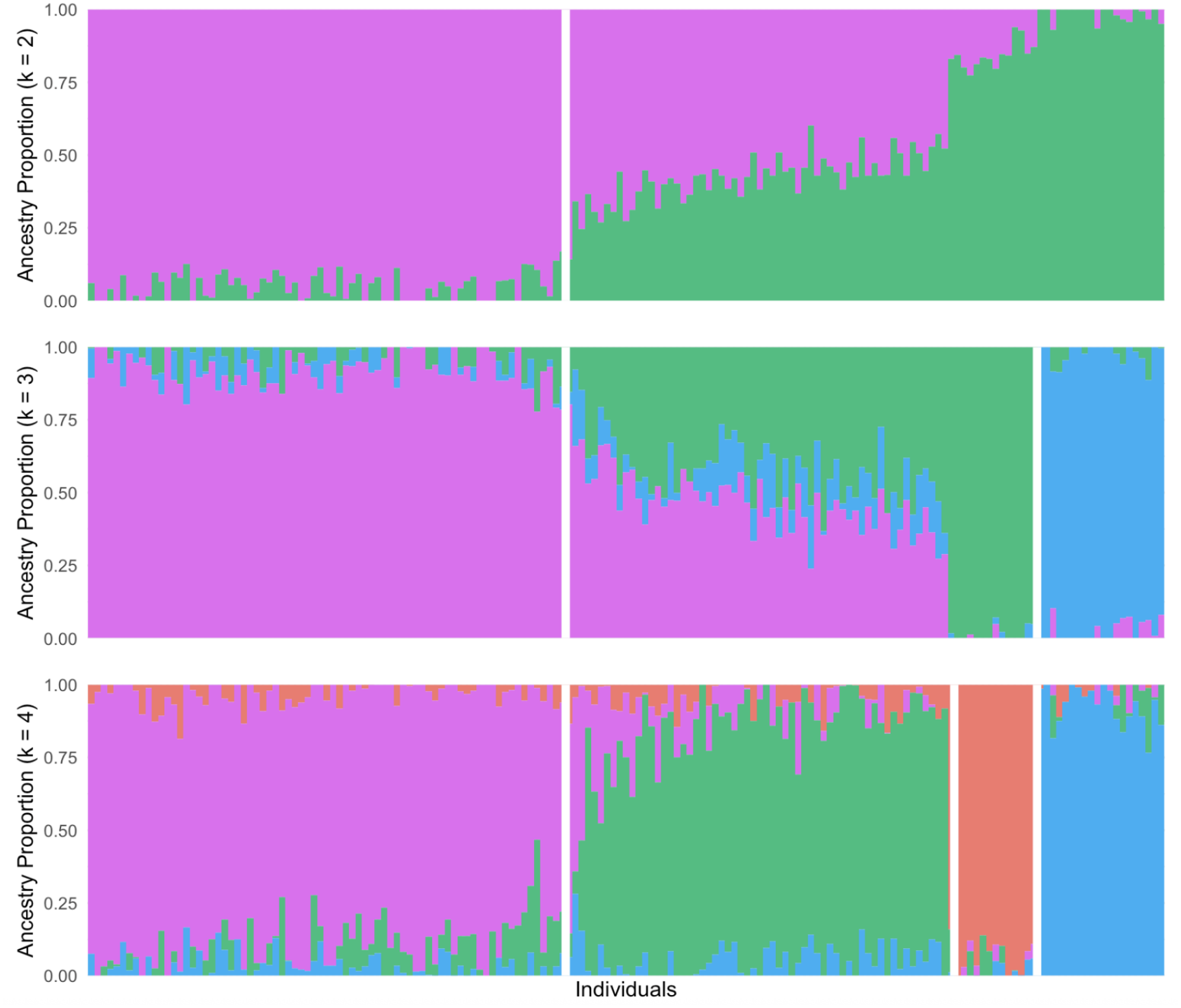
Plots depicting ancestry clusters assigned by snmf for the *G. demissa* dataset at k = 2, k = 3, and k =4 ancestral populations, respectively. Bars on the plot represent individuals, and the color within each bar represents assigned cluster identity for a given proportion of each individual’s SNPs. Individuals are arranged south-north from left to right, with the leftmost individuals sampled from Ponce Inlet, FL and the rightmost individuals sampled from Antigonish, NS. White lines indicate boundaries of each major cluster, denoting Cape Hatteras at *k = 2,* the Bay of Fundy at *k = 3*, and Cape Cod at *k = 4*.

### Part F. Alternative parameterizations for ocean model

Below are posterior distributions for contiguous Lineage Source Region descendants at each sampled location (from south to north on the Atlantic coast) for 3 different pelagic larval durations (10, 18, or 26 days).

**Supplementary Table 7:** Posterior probabilities for genomic ancestry from each lineage source region (LSR) assuming a pelagic larval duration (PLD) of 10 days.

|  | pop | region | lat | long | lsr0 | lsr1 | lsr2 | lsr3 | lsr4 | lsr5 | lsr6 | lsr7 | lsr8 | lsr9 | outside |
|---|---|---|---|---|---|---|---|---|---|---|---|---|---|---|---|
| 19 | 20 | south | 29.134 | -80.955 | 0.00E+00 | 0 | 8.77E-01 | 0.00E+00 | 0.00E+00 | 0 | 0 | 0 | 0 | 0 | 0.1228 |
| 18 | 19 | south | 29.833 | -81.325 | 0.00E+00 | 0 | 8.77E-01 | 0.00E+00 | 0.00E+00 | 0 | 0 | 0 | 0 | 0 | 0.1228 |
| 17 | 18 | south | 31.392 | -81.271 | 0.00E+00 | 0 | 8.77E-01 | 0.00E+00 | 0.00E+00 | 0 | 0 | 0 | 0 | 0 | 0.12276667 |
| 16 | 17 | south | 31.083 | -81.425 | 0.00E+00 | 0 | 8.77E-01 | 0.00E+00 | 0.00E+00 | 0 | 0 | 0 | 0 | 0 | 0.1228 |
| 15 | 16 | south | 32.038 | -81.048 | 0.00E+00 | 0 | 8.77E-01 | 0.00E+00 | 0.00E+00 | 0 | 0 | 0 | 0 | 0 | 0.1228 |
| 14 | 15 | south | 32.762 | -79.967 | 6.67E-05 | 0 | 8.77E-01 | 0.00E+00 | 0.00E+00 | 0 | 0 | 0 | 0 | 0 | 0.1228 |
| 13 | 14 | south | 34.175 | -77.932 | 1.43E-03 | 0 | 8.75E-01 | 6.67E-05 | 3.33E-05 | 0 | 0 | 0 | 0 | 0.00043333 | 0.12256667 |
| 12 | 13 | south | 34.702 | -76.751 | 1.14E-02 | 0 | 8.63E-01 | 8.67E-04 | 6.67E-05 | 0 | 0 | 0 | 0 | 0.00323333 | 0.1212 |
| 11 | 11 | mab | 37.307 | -76.411 | 9.82E-01 | 0 | 0.00E+00 | 6.67E-05 | 0.00E+00 | 0 | 0 | 0 | 0 | 0.00026667 | 0.0173 |
| 10 | 10 | mab | 38.041 | -75.363 | 2.43E-01 | 0 | 0.00E+00 | 6.33E-01 | 5.92E-02 | 0 | 0 | 0.01956667 | 0 | 0.00606667 | 0.0395 |
| 9 | 9 | mab | 39.234 | -75.031 | 1.30E-01 | 0 | 0.00E+00 | 7.34E-01 | 6.54E-02 | 0 | 0 | 0.0245 | 0 | 0.00293333 | 0.04333333 |
| 8 | 8 | mab | 39.555 | -74.363 | 7.25E-02 | 0 | 3.33E-05 | 7.83E-01 | 7.51E-02 | 0 | 0 | 0.02553333 | 0 | 0.00183333 | 0.04196667 |
| 7 | 7 | mab | 40.617 | -73.504 | 4.53E-03 | 0 | 0.00E+00 | 8.43E-01 | 8.45E-02 | 0 | 0 | 0.0273 | 0 | 0.0001 | 0.04003333 |
| 6 | 6 | mab | 40.897 | -73.434 | 0.00E+00 | 0 | 0.00E+00 | 8.59E-01 | 7.48E-02 | 0 | 0 | 0.02733333 | 0 | 0 | 0.03893333 |
| 5 | 5 | mab | 41.576 | -70.639 | 0.00E+00 | 0 | 0.00E+00 | 7.08E-01 | 2.21E-01 | 0 | 0 | 0.0226 | 0 | 0 | 0.03503333 |
| 4 | 4 | gom | 42.759 | -70.891 | 0.00E+00 | 0 | 0.00E+00 | 0.00E+00 | 9.76E-01 | 0 | 0 | 0 | 0 | 0 | 0.0237 |
| 3 | 3 | gom | 44.014 | -69.538 | 0.00E+00 | 0 | 0.00E+00 | 0.00E+00 | 9.75E-01 | 0 | 0 | 0 | 0 | 0 | 0.02363333 |
| 2 | 2 | maritimes | 44.651 | -63.423 | 0.00E+00 | 0.8328667 | 0.00E+00 | 0.00E+00 | 0.00E+00 | 0 | 0.00133333 | 0 | 0 | 0 | 0.165 |
| 1 | 1 | maritimes | 45.668 | -61.878 | 0.00E+00 | 0.8338 | 0.00E+00 | 0.00E+00 | 0.00E+00 | 0 | 0 | 0 | 0 | 0 | 0.1662 |

**Supplementary Table 8:** Posterior probabilities for genomic ancestry from each lineage source region (LSR) assuming a pelagic larval duration (PLD) of 18 days.

|  | pop | region | lat | long | lsr0 | lsr1 | lsr2 | lsr3 | lsr4 | lsr5 | lsr6 | outside |  |
|---|---|---|---|---|---|---|---|---|---|---|---|---|---|
| 19 | 20 | south | 29.134 | -80.955 | 0.00E+00 | 0 | 0.8934 | 0 | 0 | 0.00E+00 | 0.00E+00 | 0.07973333 | 9.73E-01 |
| 18 | 19 | south | 29.833 | -81.325 | 0.00E+00 | 0 | 0.9182 | 0 | 0 | 0.00E+00 | 0.00E+00 | 0.0818 | 1.00E+00 |
| 17 | 18 | south | 31.392 | -81.271 | 3.33E-05 | 0 | 0.9142 | 0 | 0 | 0.00E+00 | 0.00E+00 | 0.08143333 | 9.96E-01 |
| 16 | 17 | south | 31.083 | -81.425 | 0.00E+00 | 0 | 0.9173667 | 0 | 0 | 0.00E+00 | 0.00E+00 | 0.0818 | 9.99E-01 |
| 15 | 16 | south | 32.038 | -81.048 | 0.00E+00 | 0 | 0.9182 | 0 | 0 | 0.00E+00 | 0.00E+00 | 0.0818 | 1.00E+00 |
| 14 | 15 | south | 32.762 | -79.967 | 0.00E+00 | 0 | 0.9181667 | 0 | 0 | 0.00E+00 | 6.67E-05 | 0.08176667 | 1.00E+00 |
| 13 | 14 | south | 34.175 | -77.932 | 2.97E-03 | 0 | 0.9149 | 0.00013333 | 0 | 0.00E+00 | 5.67E-04 | 0.08143333 | 1.00E+00 |
| 12 | 13 | south | 34.702 | -76.751 | 1.70E-02 | 0 | 0.8879667 | 0.0013 | 0.0001 | 6.67E-05 | 5.43E-03 | 0.07853333 | 9.90E-01 |
| 11 | 11 | mab | 37.307 | -76.411 | 9.98E-01 | 0 | 0 | 0.0001 | 0 | 3.33E-05 | 2.00E-04 | 0.0019 | 1.00E+00 |
| 10 | 10 | mab | 38.041 | -75.363 | 1.59E-01 | 0.00503333 | 0 | 0.66206667 | 0.1309 | 1.40E-02 | 2.33E-03 | 0.02656667 | 1.00E+00 |
| 9 | 9 | mab | 39.234 | -75.031 | 8.71E-02 | 0.00563333 | 0 | 0.72003333 | 0.1413333 | 1.60E-02 | 1.40E-03 | 0.0285 | 1.00E+00 |
| 8 | 8 | mab | 39.555 | -74.363 | 5.18E-02 | 0.0055 | 0 | 0.73883333 | 0.1490667 | 1.64E-02 | 5.00E-04 | 0.029 | 9.91E-01 |
| 7 | 7 | mab | 40.617 | -73.504 | 3.37E-03 | 0.00676667 | 0 | 0.77856667 | 0.1602667 | 1.74E-02 | 1.33E-04 | 0.02946667 | 9.96E-01 |
| 6 | 6 | mab | 40.897 | -73.434 | 0.00E+00 | 0.00616667 | 0 | 0.79126667 | 0.1555333 | 1.75E-02 | 0.00E+00 | 0.02933333 | 1.00E+00 |
| 5 | 5 | mab | 41.576 | -70.639 | 3.33E-05 | 0.01623333 | 0 | 0.60593333 | 0.2698 | 1.25E-02 | 0.00E+00 | 0.02673333 | 9.31E-01 |
| 4 | 4 | gom | 42.759 | -70.891 | 0.00E+00 | 0.07326667 | 0 | 0 | 0.8986667 | 0.00E+00 | 0.00E+00 | 0.02533333 | 9.97E-01 |
| 3 | 3 | gom | 44.014 | -69.538 | 0.00E+00 | 0.07293333 | 0 | 0 | 0.8963 | 0.00E+00 | 0.00E+00 | 0.025 | 9.94E-01 |
| 2 | 2 | maritimes | 44.651 | -63.423 | 0.00E+00 | 0.94816667 | 0 | 0 | 0 | 0.00E+00 | 0.00E+00 | 0.03946667 | 9.88E-01 |
| 1 | 1 | maritimes | 45.668 | -61.878 | 0.00E+00 | 0.95993333 | 0 | 0 | 0 | 0.00E+00 | 0.00E+00 | 0.04006667 | 1.00E+00 |

**Supplementary Table 9:** Posterior probabilities for genomic ancestry from each lineage source region (LSR) assuming a pelagic larval duration (PLD) of 26 days.

|  | pop | region | lat | long | lsr0 | lsr1 | lsr2 | lsr3 | lsr4 | lsr5 | lsr6 | outside |  |
|---|---|---|---|---|---|---|---|---|---|---|---|---|---|
| 19 | 20 | south | 29.134 | -80.955 | 0.00E+00 | 0.00E+00 | 7.53E-01 | 0.00E+00 | 0.00E+00 | 0.00E+00 | 0.00E+00 | 0.06683333 | 8.20E-01 |
| 18 | 19 | south | 29.833 | -81.325 | 0.00E+00 | 0.00E+00 | 9.18E-01 | 0.00E+00 | 0.00E+00 | 0.00E+00 | 0.00E+00 | 0.0816 | 1.00E+00 |
| 17 | 18 | south | 31.392 | -81.271 | 0.00E+00 | 0.00E+00 | 8.64E-01 | 0.00E+00 | 0.00E+00 | 0.00E+00 | 0.00E+00 | 0.0764 | 9.40E-01 |
| 16 | 17 | south | 31.083 | -81.425 | 0.00E+00 | 0.00E+00 | 8.87E-01 | 0.00E+00 | 0.00E+00 | 0.00E+00 | 0.00E+00 | 0.07906667 | 9.66E-01 |
| 15 | 16 | south | 32.038 | -81.048 | 0.00E+00 | 0.00E+00 | 9.18E-01 | 0.00E+00 | 0.00E+00 | 0.00E+00 | 0.00E+00 | 0.0816 | 1.00E+00 |
| 14 | 15 | south | 32.762 | -79.967 | 3.33E-05 | 0.00E+00 | 9.18E-01 | 0.00E+00 | 0.00E+00 | 0.00E+00 | 0.00E+00 | 0.0816 | 1.00E+00 |
| 13 | 14 | south | 34.175 | -77.932 | 2.20E-03 | 6.67E-05 | 9.15E-01 | 2.67E-04 | 0.00E+00 | 0.00E+00 | 5.33E-04 | 0.08146667 | 1.00E+00 |
| 12 | 13 | south | 34.702 | -76.751 | 1.62E-02 | 1.67E-04 | 8.46E-01 | 1.67E-03 | 3.67E-04 | 3.33E-05 | 5.83E-03 | 0.0753 | 9.46E-01 |
| 11 | 11 | mab | 37.307 | -76.411 | 9.26E-01 | 2.80E-03 | 0.00E+00 | 4.65E-02 | 5.57E-03 | 5.33E-04 | 2.33E-04 | 0.01826667 | 1.00E+00 |
| 10 | 10 | mab | 38.041 | -75.363 | 1.01E-01 | 6.72E-02 | 1.33E-04 | 6.66E-01 | 1.12E-01 | 5.87E-03 | 1.03E-03 | 0.04656667 | 1.00E+00 |
| 9 | 9 | mab | 39.234 | -75.031 | 5.90E-02 | 7.07E-02 | 3.33E-05 | 6.99E-01 | 1.18E-01 | 6.07E-03 | 4.33E-04 | 0.0474 | 1.00E+00 |
| 8 | 8 | mab | 39.555 | -74.363 | 3.80E-02 | 7.22E-02 | 0.00E+00 | 6.75E-01 | 1.16E-01 | 5.93E-03 | 4.00E-04 | 0.04563333 | 9.52E-01 |
| 7 | 7 | mab | 40.617 | -73.504 | 3.60E-03 | 8.33E-02 | 0.00E+00 | 7.15E-01 | 1.26E-01 | 6.20E-03 | 3.33E-05 | 0.04783333 | 9.82E-01 |
| 6 | 6 | mab | 40.897 | -73.434 | 1.00E-04 | 8.23E-02 | 0.00E+00 | 7.35E-01 | 1.27E-01 | 6.27E-03 | 0.00E+00 | 0.0493 | 1.00E+00 |
| 5 | 5 | mab | 41.576 | -70.639 | 1.33E-04 | 2.20E-01 | 0.00E+00 | 5.00E-01 | 1.73E-01 | 4.57E-03 | 0.00E+00 | 0.0538 | 9.52E-01 |
| 4 | 4 | gom | 42.759 | -70.891 | 0.00E+00 | 6.16E-01 | 0.00E+00 | 4.33E-04 | 3.02E-01 | 0.00E+00 | 0.00E+00 | 0.07466667 | 9.93E-01 |
| 3 | 3 | gom | 44.014 | -69.538 | 0.00E+00 | 6.05E-01 | 0.00E+00 | 4.00E-04 | 2.94E-01 | 0.00E+00 | 0.00E+00 | 0.07356667 | 9.73E-01 |
| 2 | 2 | maritimes | 44.651 | -63.423 | 0.00E+00 | 8.94E-01 | 0.00E+00 | 0.00E+00 | 3.33E-05 | 0.00E+00 | 0.00E+00 | 0.06043333 | 9.55E-01 |
| 1 | 1 | maritimes | 45.668 | -61.878 | 0.00E+00 | 9.37E-01 | 0.00E+00 | 0.00E+00 | 0.00E+00 | 0.00E+00 | 0.00E+00 | 0.06326667 | 1.00E+00 |

### Part G: Redundancy analysis

**Supplementary Figure 17.**
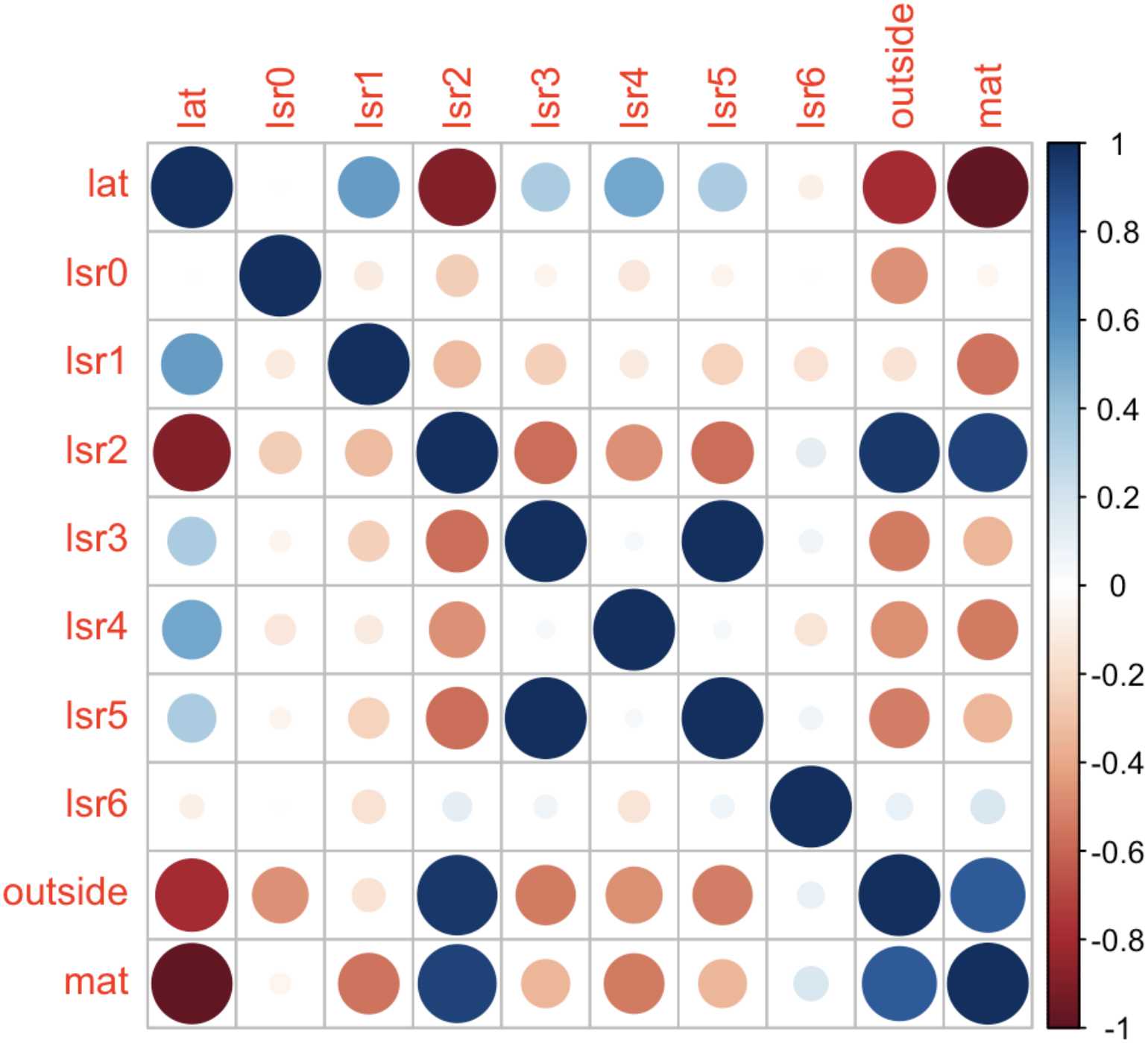
A heatmap displaying correlation coefficients between parameters included in different RDA models, where “lat” stands for latitude, “lsr” entries are the proportion of each sample predicted to descend from a given LSR, “outside” is the proportion of each sample predicted to descend from lineages outside of a given LSR, and “mat” stands for mean annual sea surface temperature. The high collinearity displayed here rendered a single model with every predictor model inappropriate.

**Supplementary Figure 18.**
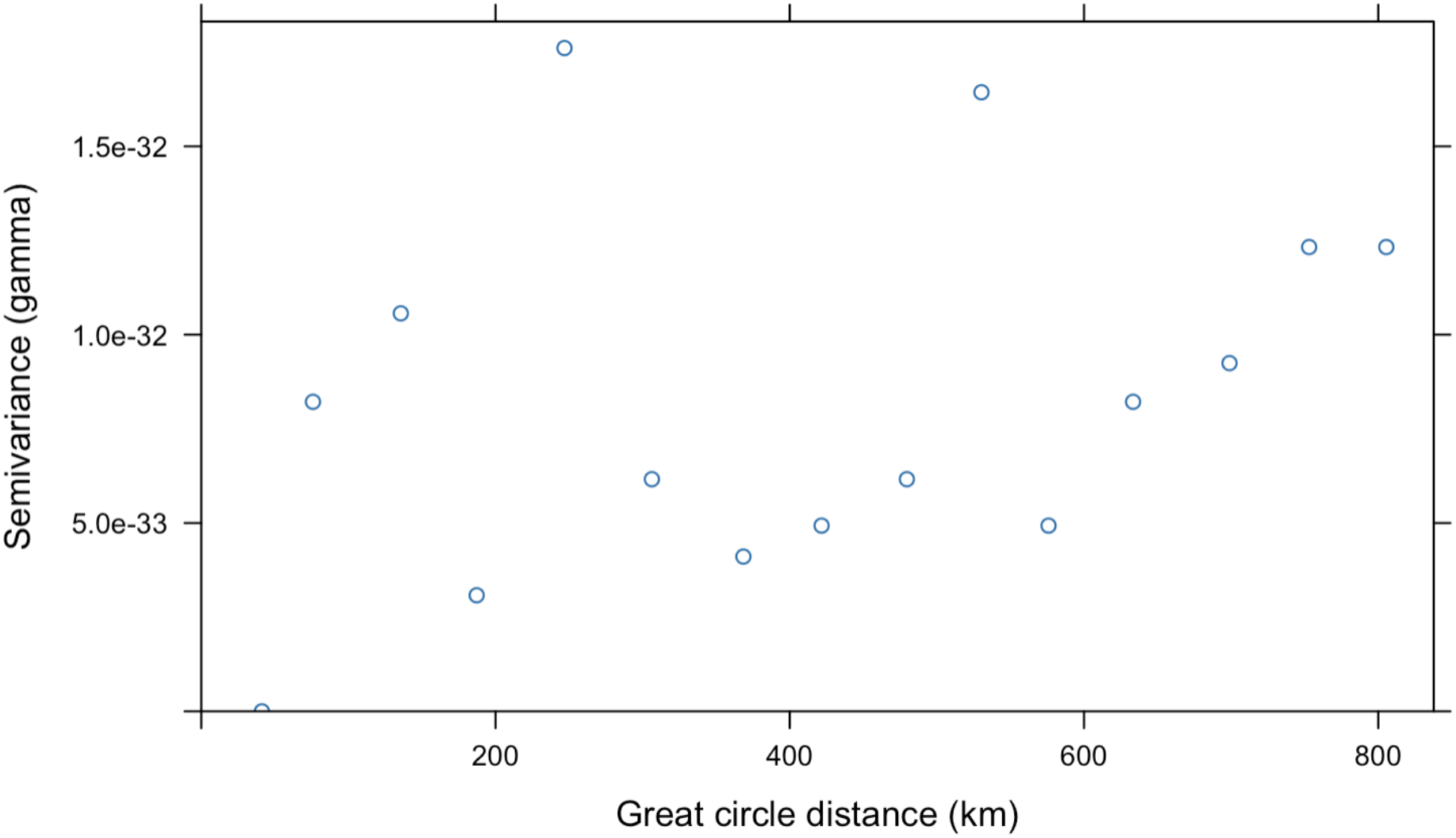
A semivariogram for the LSR RDA, illustrating semivariance of RDA1 residuals for each sampling location as a function of great circle distances between locations, is displayed above. This lack of spatial structure in residual semivariance indicates that the LSR model accounts for spatial variation in genomic data, and that variation unaccounted for by the model is likely random.

### Part H: Measuring goodness of fit relative to isolation by distance

To quantitatively assess the performance of ocean current modeling against empirical population genetic results, as well as the population genetic results against a null model of isolation by distance, we chose Hellinger distances as a measure of dissimilarity in probability distributions. Briefly, for each location *i* we have an empirical proportion of ancestry from K lineages, and a predicted ancestry of K* types from the ocean current model. We address the issue when K ≠ K* later, but the distance is then measured as

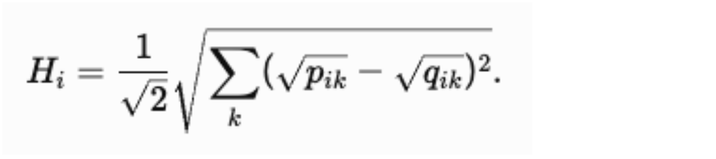

With this formulation, the Hellinger distance *H_i_* ranged from 0 to 1 for each location; we then take the root mean square deviation across *i* locations to summarize matrix correspondence across spatially autocorrelated locations. For the empirical population genetic data, *K* is determined by minimizing cross-entropy for sNMF estimates of ancestry (see Methods).

#### Isolation by distance models

We developed a spatially explicit model of symmetric stepping-stone isolation by distance using SLiM (Heller ref). We recognize from the empirical Results that the maximum F_ST_ among sites south of Cape Cod, which have not experienced bottlenecks associated with the last glacial maximum, is ∼0.03 across a spatial range of ∼1870 km. The maximum Fst is ∼0.10 across the entire range of *G. demissa*, a spatial range of ∼2840km.

Our SLiM simulation was then set to simulate a 6Mbp chromosome evolving with a mutation rate (and recombination rate) of 1e-8, with all mutations having no fitness effect. Sites have population sizes of 1000 and migration rate m0 tuned to 3 values [0.01, 0.025, 0.05] which lead to output F_ST_ values over the spatial domain of *G. demissa* approximating those noted above plus an intermediate value. We note that F_ST_ values themselves are not directly indicative of migration patterns for a number of reasons (16). Migration was entirely symmetric and distance-based. Simulations run for 10^4^ generations.

Resultant VCF files are converted to .geno format for use in the LEA package and snmf analysis; a subset of SNPs are randomly selected to match the variable sites in our empirical data, and 10 individuals are sampled from each of the simulated populations. These simulated IBD replicates are then analyzed for K values of ancestry as with the empirical data.

Recognizing that in a pattern of isolation by distance for any value of K, one ancestry type will be dominant at either end of the distribution with other types gaining dominance in intermediate sites, each output Q’ matrix was re-sorted by the spatial dominance of inferred ancestry type from south-to-north so that all are directly comparable. The same re-sorting is performed on the empirical Q matrix from the population genetic data. An example of spatial variation in empirical (left) as well as simulated ancestry (right) across the spatial range of our study is shown below.

**Supplementary Figure 19:**
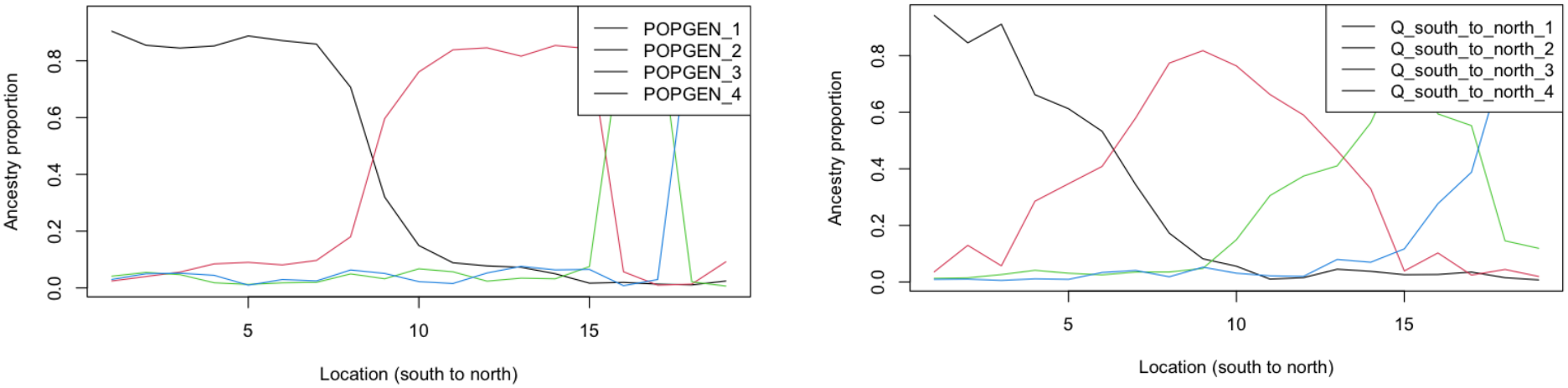
Density plots representing proportion of genomic ancestry attributable to each of four clusters. The left and right panels representsancestry coefficients before and after north-south reorganization, respectively.

#### SLiM code

The code is written to be called from command-line interface providing the value of M0 to parameterize the simulation as above.

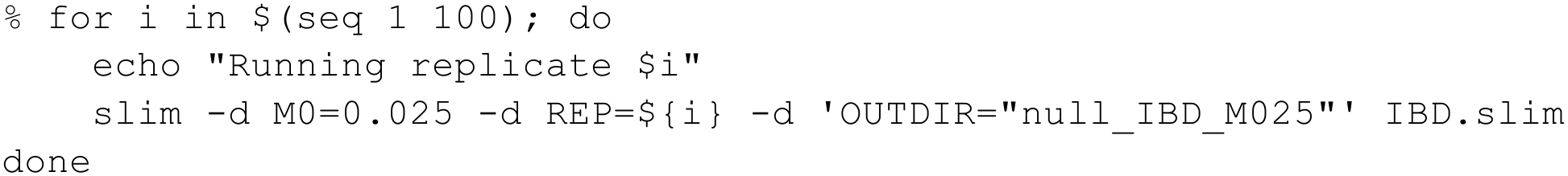

The code includes pairwise spatial distances in kilometers that are generated from the latitude-longitude of each location. As written, the code includes Lake Worth, Florida but does not include Roanoke Sound, NC because of the near-complete introgression by the Gulf ribbed mussel.

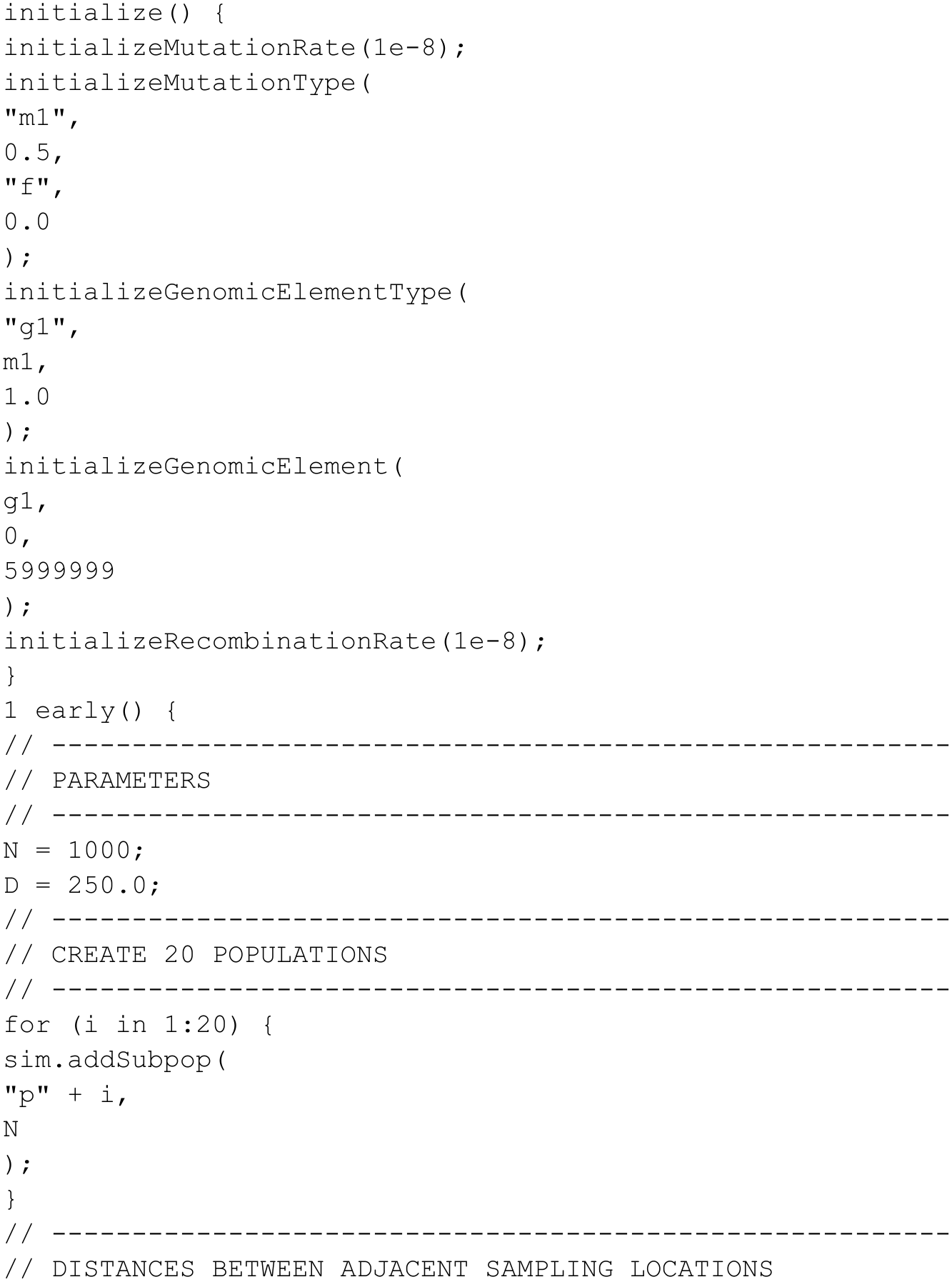
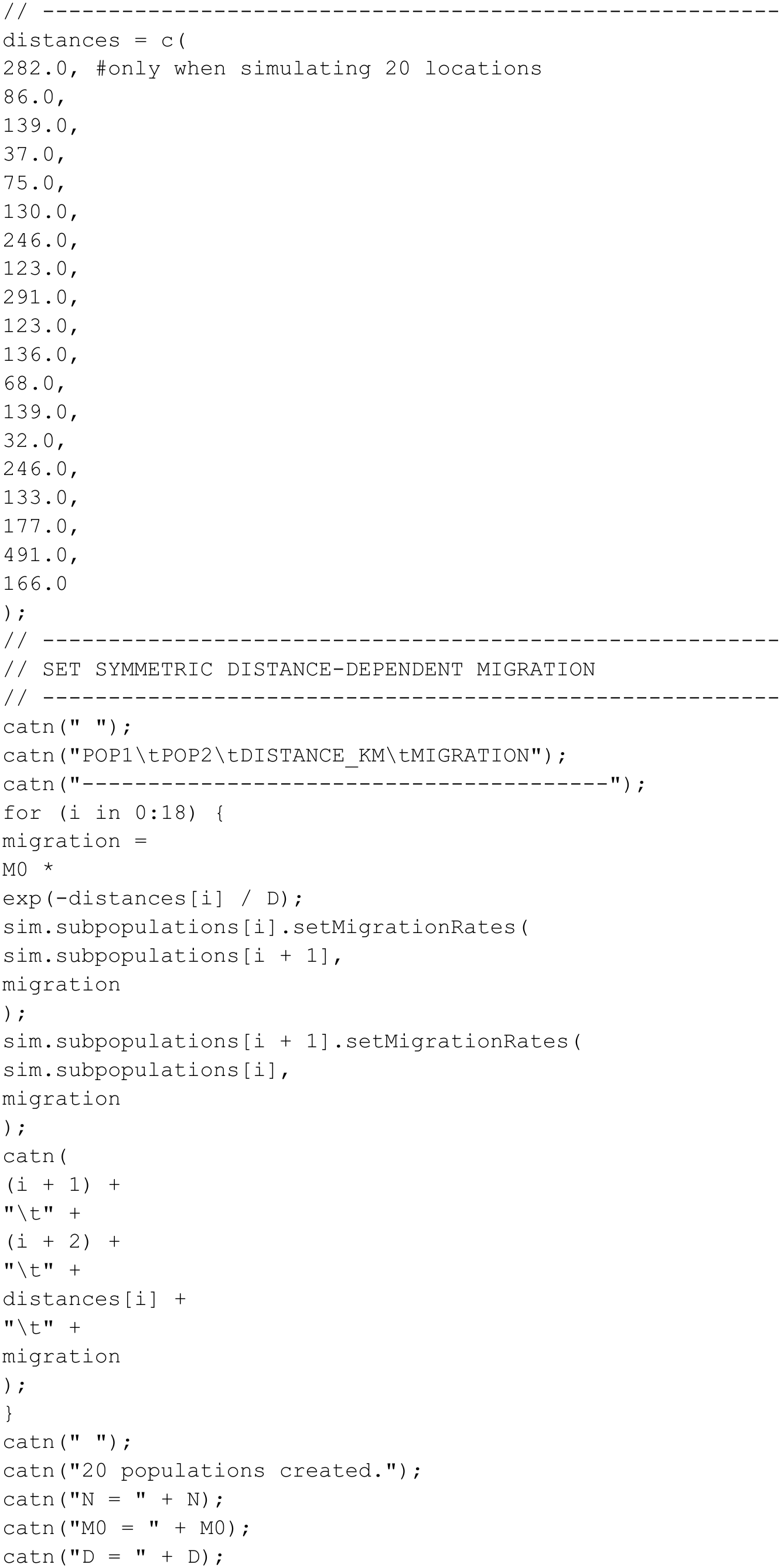
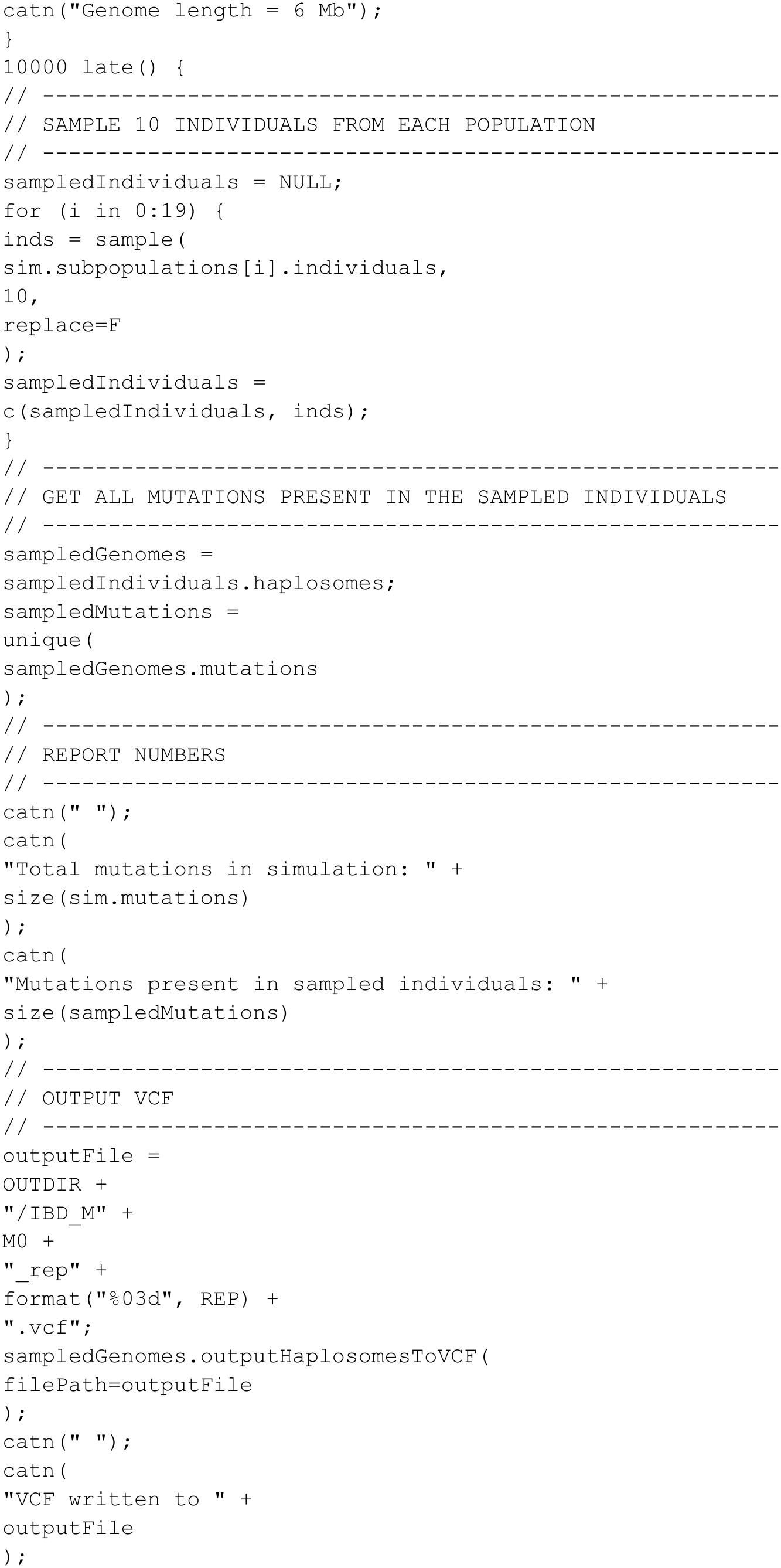
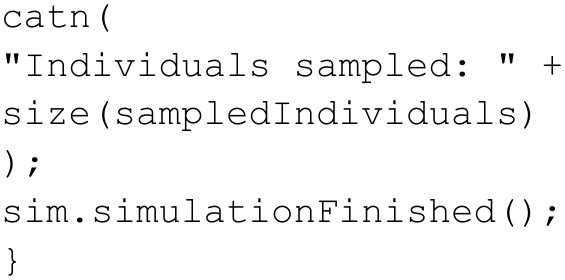

#### Merging predicted types from physical ocean model

To calculate the Hellinger distance between any two complex distributions, they should have the same number *k* of factors being compared. As the physical ocean model used herein is based purely on physical dispersal of simulated particles which are then back-tracked to their points of origin, the number of Lineage Source Regions (LSRs) includes some small regions with minor contributions from a distinct LSR as well as some particles that are retained within a broader region that did not originate from the upstream LSR *sensu stricto* but from elsewhere (typically within the same LSR-dominated region).

As the identity of types in the physical ocean model have no evolutionary relatedness to one another - as with the infinite alleles model in population genetics - we simplify the results of the full ocean current models to reflect the dominant inferred types. LSRs that contribute less than 5% to the overall predicted diversity, and the contribution of diversity not directly descended from an LSR, are excluded. This approach still permits the remaining LSR distributions to predict the ancestry for 75.9% (PLD26) to 89.2% (PLD10) of the sampled range of *G. demissa*.

Row frequencies are normalized to sum to 1 and the forecasts of ancestry descendant from each LSR are established by fixing the northern/southern-most arrays to match the northern/southern-most of comparison arrays (as in the IBD simulations which by definition have distinct nearly-fixed types at each end of the domain), and then summing the interior LSR arrays by their regional predictive affinity (i.e. collapsing neighboring predictions). The ocean current model only predicts based on areas of high retention of local diversity, but stochastic events of gene flow can always replace the diversity in any region with that of another – thus the merging of diversity allows us to compare the optimal K=4 from the empirical data with 4 distributions in the ocean model that reflect most of the predicted diversity based on physics alone.

#### Summary of Results

Given 3 different parameters for the physical ocean model to assess the stability of prediction across distinct assumptions of larval dispersal, and 3 different parameters for simulated migration tuned to observed and intermediate F_ST_ across the distributional range, we summarize the value of the physical model over a null expectation of IBD by reporting the percentile of Hellinger distance from genetic results to each physical model relative to the distance to the distribution of IBD results. In most comparisons, the physical model has higher concordance with genomic structure than 90% of IBD simulations.

**Supplementary Table 10:** Fit of the LSR model to empirical data as represented by the percentage of isolation-by-distance (IBD) simulations for which its Hellinger distance is lower. Higher percentile values indicate improved performance over the IBD null.

|  | M<sub>0</sub> = 0.01 | M<sub>0</sub> = 0.025 | M<sub>0</sub> = 0.05 |
|---|---|---|---|
| PLD 10 days | 94 | 89 | 91 |
| PLD 18 days | 94 | 89 | 91 |
| PLD 26 days | 87 | 76 | 78 |

